# An acoustic platform for single-cell, high-throughput measurements of the viscoelastic properties of cells

**DOI:** 10.1101/2020.09.07.286898

**Authors:** Valentin Romanov, Giulia Silvani, Huiyu Zhu, Charles D Cox, Boris Martinac

## Abstract

Cellular processes including adhesion, migration and differentiation are governed by the distinct mechanical properties of each cell. Importantly, the mechanical properties of individual cells can vary depending on local physical and biochemical cues in a time-dependent manner resulting in significant inter-cell heterogeneity. While several different methods have been developed to interrogate the mechanical properties of single cells, throughput to capture this heterogeneity remains an issue. While new high-throughput techniques are slowly emerging, they are primarily aimed at characterizing cells in suspension, whereas high-throughput measurements of adherent cells have proven to be more challenging. Here, we demonstrate single-cell, high-throughput characterization of adherent cells using acoustic force spectroscopy. We demonstrate that cells undergo marked changes in viscoelasticity as a function of temperature, the measurements of which are facilitated by a closed microfluidic culturing environment that can rapidly change temperature between 21 °C and 37 °C. In addition, we show quantitative differences in cells exposed to different pharmacological treatments specifically targeting the membrane-cytoskeleton interface. Further, we utilize the high-throughput format of the AFS to rapidly probe, in excess of 1000 cells, three different cell-lines expressing different levels of a mechanosensitive protein, Piezo1, demonstrating the ability to differentiate between cells based on protein expression levels.

## INTRODUCTION

The response of livings cells to externally applied loads is incredibly diverse and reflective of their organization, function and location. How cells respond to these external cues will inform their behavior, forming a complex cascade of cellular signaling events consisting of the conversion of physical forces into biochemical cues referred to as mechanotransduction^1^. The alteration of cellular characteristics as part of this signaling cascade has been linked to the onset and progression of human disease^2^, including Sickle cell anemia, arthritis and cancer^3^. Mechanical cues such as compression have been shown to shift the behavior of cancer cells, promoting a more invasive phenotype^4^. Force directionality plays an important part in mechanotransduction and for reproducing normal phenotypes when studying cellular mechanics. For instance, compressive forces are less representative of the environment experienced by those cells found in the heart, gut and the lung, that are subject to stretching forces^5^. As the force application and directionality or environmental conditions change, so do the mechanical properties of cells, including changes in stiffness and/or fluidity. As such, understanding how the mechanical and viscoelastic properties of cells vary as a function of force, environment and chemical and biological cues is of critical importance.

Many techniques have been developed to study the mechanical properties of cells^6,7^, these include magnetic twisting cytometry (MCT)^8^, atomic force microscopy (AFM)^9^, optical tweezers (OT)^2^, micropipette aspiration^10^ and particle tracking microrheology (PTMR)^11^. Recently, acoustic force spectroscopy (AFS) has emerged as an alternative method for providing high-throughput tracking at single-cell resolution^12,13^. Conceptually, the AFS is similar in operation to Optical Tweezers. When optically trapped, membrane-attached single particles (a.k.a handles) are pulled away from the cell surface, a thin tether is formed. Likewise, the AFS utilizes acoustically driven, membrane-attached particles as “handles” for pulling tethers from 30 to 50 cells simultaneously^13^. As such, the AFS has great potential as a diagnostic tool for characterizing the mechanical properties of adherent cells. Previously, the AFS was used to characterize the mechanical properties of red blood cells exposed to a variety of chemical treatments, including changes to the lipid membrane by vesicle fusion^13^. Recently, the AFS was used to measure the complex shear modulus of human umbilical vein endothelial cells^14^.

Numerous reports have been published looking at the effect of either cytoskeletal or plasma membrane modifications on the viscoelastic properties of cells^15^. For instance, optical tweezers have been used to show cell stiffness and fluidity modulation based on enrichment or depletion of cholesterol^16^. Changes to protein expression, whether through over-expression of lamin A/C, an intermediate filament protein localized in the nuclear envelope^17^ or the knockout of vinculin, a protein of the focal adhesion complex^18^, have been shown to have a measurable effect on the viscoelastic properties of cells. Here, we utilize the AFS to demonstrate high-throughput, single-cell characterization of Human Embryonic Kidney (HEK) cells, capturing the heterogeneity inherent to all cells. We demonstrate that temperature has a dramatic impact on mechanical cell properties and that the AFS can be used to rapidly modulate cellular temperature within the microfluidic environment. We probe the viscoelastic properties of HEK293T cells with pharmacological treatments, showing that the AFS can accurately capture the mechanical properties at the membrane-cytoskeletal interface. Further, we demonstrate that the AFS can be used to rapidly, in high-throughput fashion, characterize the mechanical properties of cells based on the expression levels of a target mechanosensitive protein, Piezo1.

## RESULTS AND DISCUSSION

### A simple workflow for culturing and probing the viscoelastic properties of cells

One of the main advantages of the AFS is the “plug-and-play” nature of the technique. Contextually, plug refers to the simple act of loading the microfluidic device (AFS chip) with cells to be analyzed (**Figure 1a**). Play, refers to the act of interfacing the AFS chip with the moving stage, at which point the system is ready (no further initiation is needed). Another advantage of the system is the enclosed, glass microfluidic channel which can be functionalized with molecules of choice. Once coated, the AFS chip is seeded with cells (**Figure 1b**) and placed directly into an incubator set to 37 °C with 5% CO_2_. Cells are allowed to adhere for a predetermined amount of time, in this case 3 hrs (**Figure 1c**). The chip is then removed from the incubator and placed onto the AFS stage, at which point the temperature within the chip is set to 37 °C. The cells are now ready for probing. Silica beads can be added either during the initial cell loading stage or after the cells have adhered to the underlaying substrate.

**Figure 1.**
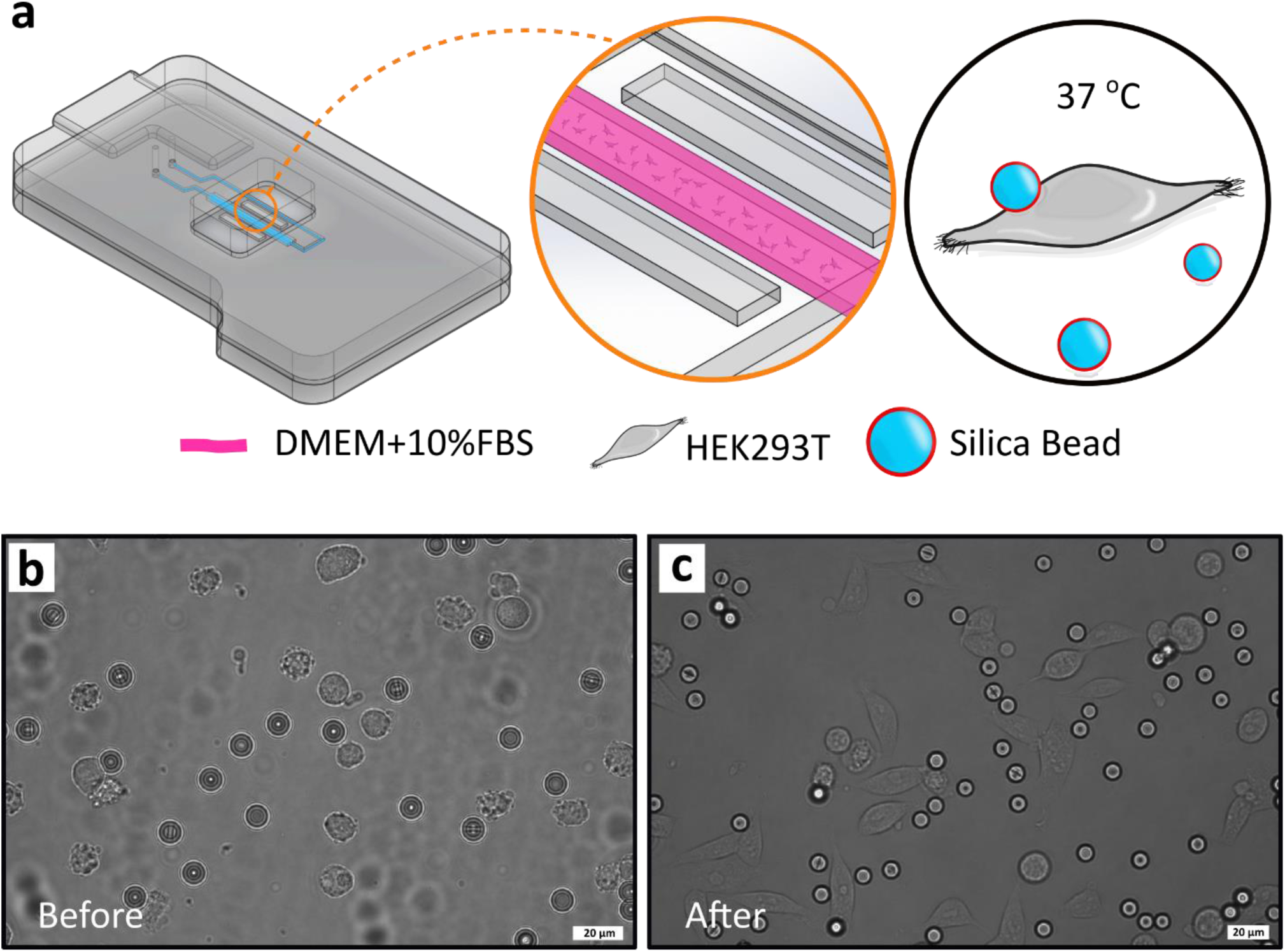
Cell culturing. **a)** A cartoon representation of the AFS chip and fluidic channel loaded with cells. HEK293T cells, together with the Silica beads are cultured directly inside the AFS device over a period of 3 hours. Channel width, length and height are 2 mm, 2 cm and 100 µm, respectively, with a total approximate volume of 4 µL. **b)** HEK293T cells with Silica beads immediately after being loaded into the microfluidic channel and **c)**, cells after 3 hours at 37 °C with 5% CO_2_.

### Methodology for force calibration and cell viscoelasticity measurements

The AFS chip is a microfluidic device with an inner internal volume of approximately 4 µL. A transparent electronic piezo element is bonded to the chip, allowing for direct visualization and tracking of objects upon application of acoustic force. Acoustic force over the height of the channel is in the form of an acoustic standing wave, generated by the transparent piezo element^19^. When set to an optimal resonance frequency, objects within the microfluidic chamber are driven towards the acoustic node (**Figure 2a**). To account for hydrodynamic drag (Methods Section) the viscosity of the sample has to be known. The viscosity of the fluid can be determined using the AFS via two approaches, 1) by taking a force balance around a sedimenting particle where upon reaching terminal velocity the viscosity can be extracted, or 2) by measuring the Mean Square Displacement of particles moving under Brownian motion and using the Einstein relation to find viscosity after experimentally determining the diffusion coefficient. Here, we use the second approach, the methodology for which is described in the Methods section (**Supplementary Figure S1**). We confirm accuracy by first measuring the viscosity of water at 21 °C with 1 µm Silica particles and find, when compared to published data^20^, µ_water_/µ_AFS_ is within 4 % (n=9). The viscosity of the media used here, DMEM+10% FBS at 37 °C is determined to be 0.74±0.05 mPa.s (n=27), and is within 4% of previously published results^21,22^. Using this viscosity value, we confirmed the quadratic force scaling vs voltage relationship for each device where each device has a unique, experimentally determined operating frequency (**Figure 2b**).

**Figure 2.**
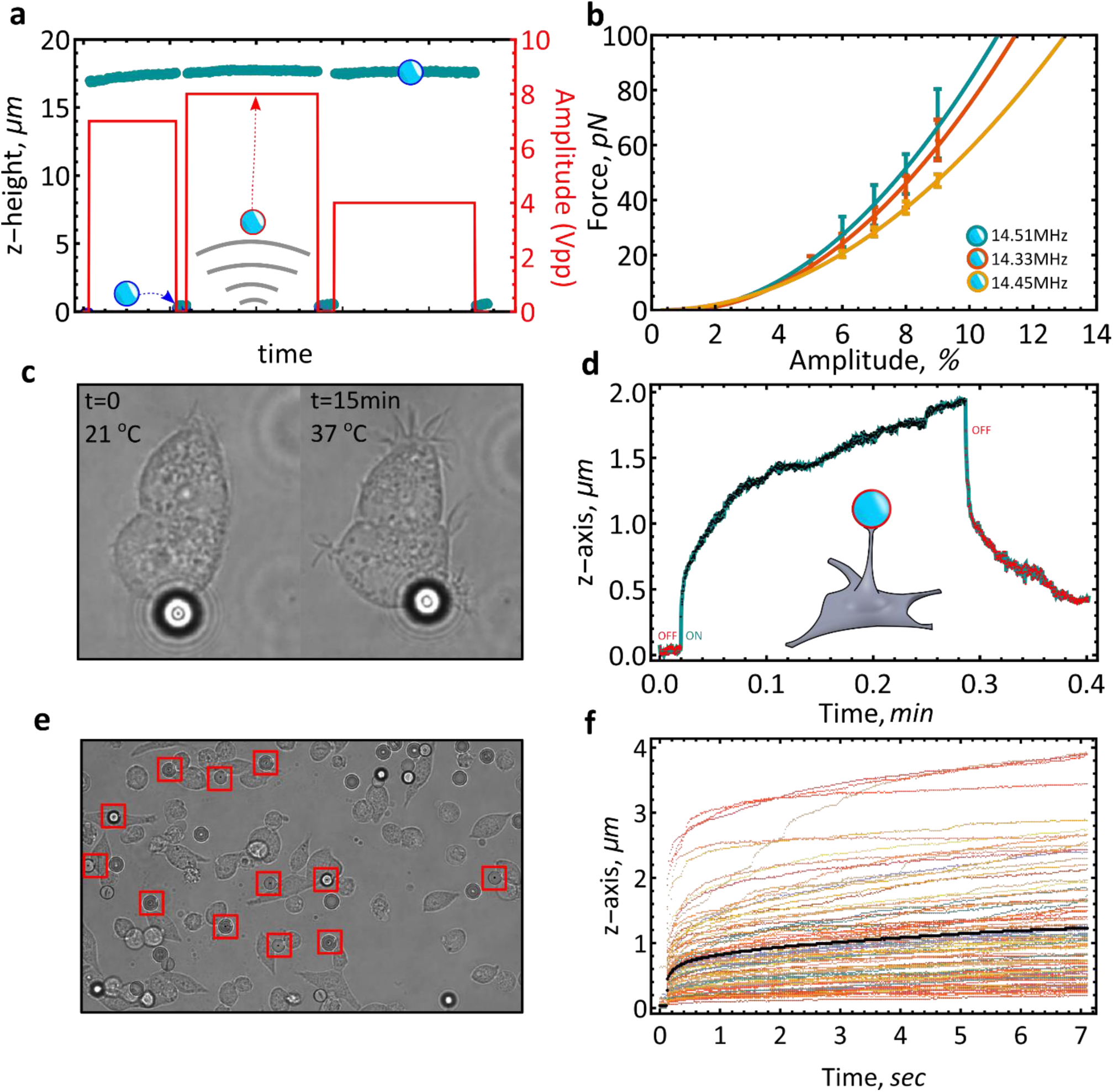
Methodology for force calibration and cell viscoelasticity measurements. **a)**, Particle movement in the x,y and z direction is tracked in real time at 60 FPS. The “particle shooting method” is used to derive force as a function of applied amplitude. **b)**, Force as a function of amplitude for 9.2 µm Silica particles in DMEM with 10 % FBS at 37 °C. Three different devices are tested. (14.51 MHz, n=10) (14.33 MHz, n=17) (14.45 MHz, n=17) (mean±S.E.M). **c)**, The fluidic chamber within the AFS device can ramp from room temperature to 37 °C in under a minute. HEK293T cells respond almost immediately to physiological temperatures by sending out filopodia. A single, attached Silica bead moves with the cell. **d)**, Demonstration of the viscoelastic behavior of HEK239T cells upon the application of a constant force step (∼29 pN). **e)**, A typical field of view for pulling experiments involving HEK293T cells. Multiple beads (red squares), attached to cells, are tracked in real-time (HEK293T cells, DMEM with 10% FBS, 9.2 µm Silica beads). **f)**, Rapid extension measurements, taken from multiple locations all over the fluidic chip, are collated into a single experiment, taken within a span of 30 minutes. In such fashion, each experiment can potentially generate hundreds of extension curves, (HEK293T cells, DMEM with 10% FBS, 9.2 µm Silica beads) (black line is the ensemble mean).

Furthermore, each device can be operated at a range of temperatures, from room temperature up to 40 °C. We probe the effectiveness of the heating element by observing the response of HEK293T cells to temperature gradients (**Figure 2c**). A typical response includes filopodia projection, cell-surface activity and bead-interaction (probing, moving). Initially at 37 °C, we then allow the chamber to cool to room temperature over a period of 30 minutes at which point the cell completely retracts all filopodia/lamellipodia. The chamber is then heated again to 37 °C and after 15 minutes the initial physiological response is restored. Using the bound silica particle, we can then probe the viscoelastic properties of the cell at any desired temperature. The extension curve characterizes the compliance of that individual cell in the form of a single creep-response (**Figure 2d**). By tracking multiple particles, it is possible to record the creep response of multiple cells upon application of an acoustic force (**Figure 2e**). Large silica particles are used to ensure optimal optical contrast against cells. We experimented with a variety of particles with different diameters and material compositions and optimized based on numerical aperture (NA) and magnification (5 µm Silica, 5 µm Polystyrene and 10 µm Silver-coated Silica particles). For the assay described here, the best contrast is given by 9.2 µm Silica particles with at least an 0.75 NA objective (**Supplementary Figure S2**). At this magnification, our field of view typically captures 10 to 40 cells (**Figure 2e**), however, depending on cell size, objective magnification and NA, several hundred cells can potentially be recorded simultaneously with potentially single-digit nanometer resolution ^13^. On average, a single 60-minute experiment yields 80 - 180 creep-force curves (**Figure 2f**) with a cell confluence of 60-80%. The simplicity and throughput of the AFS overcomes some of the traditional limitations of the more established techniques^23^.

### Validating force linearity and measurement consistency

One of the primary advantages of the AFS is real-time, high-throughput tracking of multiple cells (**Figure 3a**). The most challenging aspect is successfully tracking particle movement through the entire range of motion (from surface to the node) within the vicinity of other objects. Using the protocol developed here, we have been able to follow multiple beads through their full range of motion without losing tracking. By analyzing the extension curves, we show that the tracking is sensitive enough to differentiate between three different force steps, each step increasing by about 30 pN (**Figure 3b**). Next, we probe the relationship between the scaling modulus and the force. When exposed to external loads cells will respond in one of two ways. First, a large external load will induce stress-stiffening and fluidization within the cell. The resultant stress-stiffening behavior is non-linear, resembling a decaying exponential function. Alternatively, at small external loads the response is linear and is independent of the applied force^24^. This has been previously demonstrated using AFM, where the elastic modulus was found to be independent of applied force^25^. By force stepping from 46 to 111 pN we show the same behavior, albeit obtained by pulling rather than pushing (**figure 3c**), confirming that the acoustic force application and the cellular response to this force is linear, at small amplitude.

**Figure 3.**
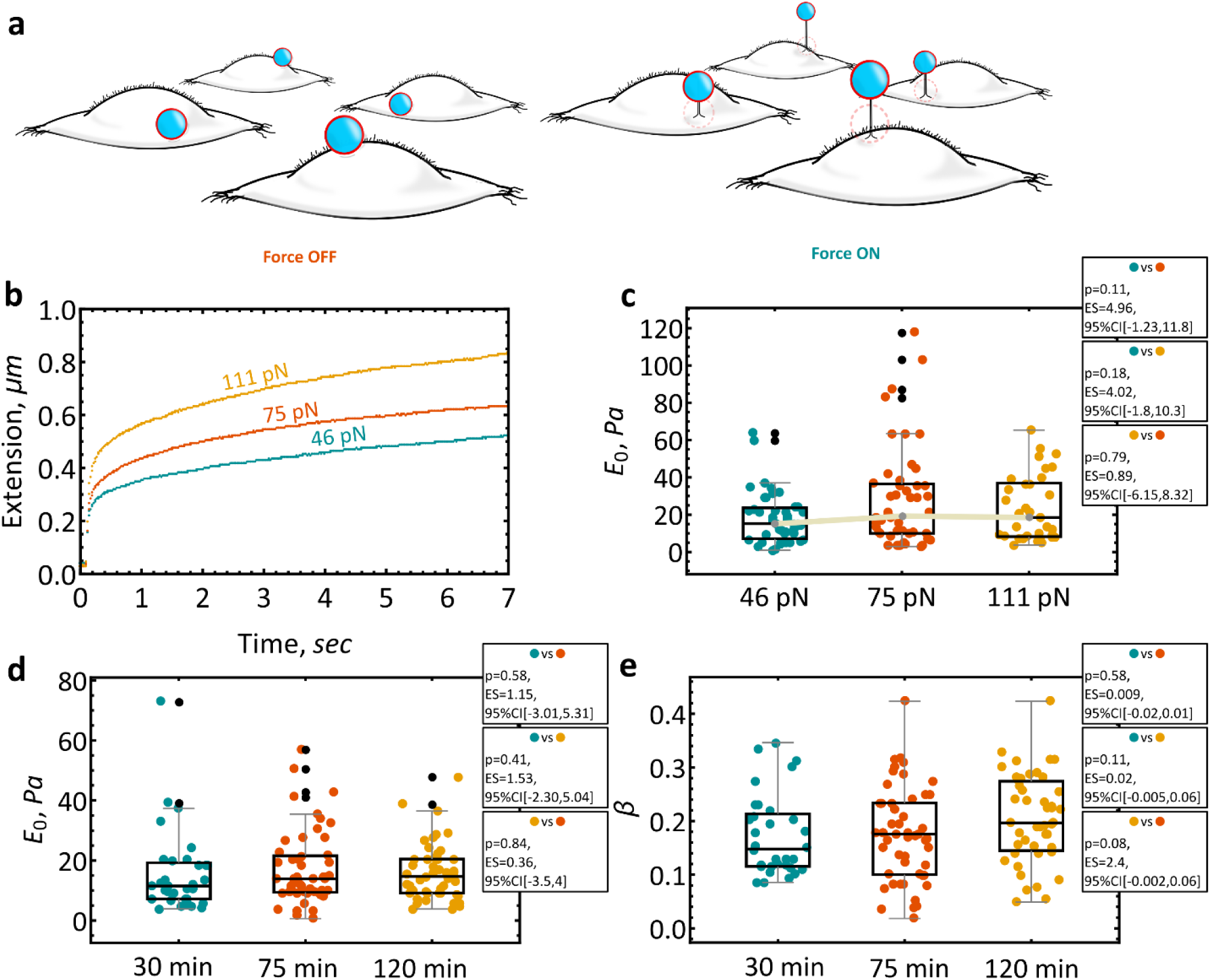
Force linearity and measurement consistency. **a)**, A cartoon representation of acoustically pulled tethers. Tether length dependence on cell type, adhesion, temperature etc. **b)**, Upon application of acoustic force, fibronectin-coated silica particles are pulled towards the acoustic node and in-turn stretch the cell (46 pN, 8 Amp%, n = 43) (75 pN, 10 Amp%, n = 43) (110 pN, 12 Amp%, n = 38) (14.33Mhz). **c)**, Force-independent linear response is confirmed by plotting the modulus scaling parameter, E_0_, as a function of force. Stress-stiffening is not observed (46 pN, E_0_ = 13.3⋇2.1 Pa, n = 43) (75 pN, E_0_ = 18.2⋇3.9 Pa, n = 33) (110 pN, E_0_ = 17.3⋇3.1 Pa, n = 48). **d)**, Cells are probed along the length of the microfluidic device. No statistically significant differences in E_0_ are found between cells measured immediately after placement onto the AFS (E_0_ = 12.1⋇2.4 Pa, n = 32), 75 minutes (E_0_ = 13.3⋇1.7 Pa, n = 51) or 120 minutes later (E_0_ = 13.6⋇1.4 Pa, n = 47). **e)**, No statistically significant differences in the power-law exponent, β, is found between cells measured directly after removal from the incubator (β = 0.16⋇0.013, n=32), 75 minutes (β = 0.17⋇0.01, n=51) or 120 minutes later (β = 0.20⋇0.01, n=47). A 58 pN (10% Amp, 14.45Mhz) distance force clamp is used for d) and e). Values are reported as the geometric mean ⋇ S.E. Black dots indicate outliers. Statistics are displayed in each figure and list the p-value, effect size (ES) and the corresponding 95% confidence interval. Statistical differences are determined using estimation statistics (see methods). Unless specified, all experiments are performed at 37 °C.

Conceptually, the AFS is similar to optical tweezers where a tether, pulled from a cell, opens the way for quantifying the viscoelastic properties of the cell. The magnitude of the viscoelastic properties is cell-type and technique dependent. For the same cell-type, these properties can vary 1000 fold, and are both technique and protocol dependent^23^. Less emphasis has been placed on understanding the variation of viscoelastic properties as a function of force directionality. By pulling membrane tethers, optical tweezers experiments have demonstrated that certain cell-types exhibit lower stiffness when compared against indentation^26^. Similar experiments have demonstrated that pulling can be used to differentiate between cell types^26^, substrate-dependent properties^27^ and drug-dependent changes^13^. While it is hard to directly compare the viscoelastic properties of cells across different techniques, force-directionality, ie pushing vs pulling, should also be considered. For example, the properties of HEK293 cells probed by a spherically-tipped or pyramidally-tipped AFM showed an elastic modulus of around 250 Pa^28,29^ or up to 2000 Pa with a conical tip^29^ whereas the modulus scaling parameter (sometimes referred to as the apparent Young’s Modulus) measured here is 10.1⋇0.57 Pa (n = 319). However, when compared to a pulling technique, the results are much closer. Using optical tweezers, the Elastic Coefficient, k_1_, of HEK293 cells has been measured at around 330 pN/ µm^30,31^ and is comparable to the k_1_ value measured using AFS here of 544⋇78 pN/µm (n = 306). Generally, the elastic modulus is about 2^27^ to 6^32^ times smaller when cells are pulled rather than indented.

The throughput is limited to cell size and microscope objective magnification where at x20 magnification around 3 to 20 cells can be probed at once. To obtain greater statistics we investigated whether the AFS chip was able to maintain cell viability over the time it takes to measure all cells inside the chamber. We utilize the Power-Law model to capture the viscoelastic properties of cells where E_0_ is the apparent elastic modulus (units of Pascal) and β, the fluidity (see Methods Section). The viscoelastic properties of a cell arise from its ability to modulate its stiffness by exhibiting both solid and viscous behavior. The cell is able to deform, flow and reorganize^33^ based on external cues, where the power-law exponent β, captures the fluidity and dissipative properties of the cell^18^. Starting close to the inlet and measuring cells in serpentine fashion, field of view, by field of view (**Supplementary Figure 3S**), we find no statistically significant differences between cells measured 30 minutes (E0 = 12.1 ⋇ 2.4 Pa) into the experiment and cells in the middle (75 min, E_0_ = 13.3 ⋇ 1.7 Pa) or those cells close to the outlet (120 min, E_0_ = 13.6 ⋇ 1.4 Pa) (**Figure 3d**). Depending on cell density, all cells within the AFS chip can be measured within 2 hrs. Cell viability depends on a number of factors including nutrients, temperature and oxygen/CO_2_ levels. For long term studies, the AFS chip can be interfaced with a pump to deliver nutrients over an extended period of time. In addition, we found no statistically significant differences in the power-law exponent (**Figure 3e**), β, the magnitude of which (∼95% CI, [0.16-0.20]) is in good agreement with previously reported values of whole-cell measurements^18,34^.

Further, we can use the results in this section to rule out localized acoustic heating as a source of potential influence on the underlaying mechanics. Optical tweezers and stretchers can induce localized heating^35^, causing structural damage to the object. Acoustic heating strongly depends on applied voltage. Figure 3d and 3e capture the viscoelastic properties of 130 cells at an applied amplitude of 10% with no statistically significant differences between cells measured at time 0 to cells measured 2 hours later, suggesting no measurable adverse effects at the structural level. Cells are continuously exposed to acoustic pressure as each force clamp is held for 8 seconds. It should be noted that acoustic pressure is applied throughout the whole chip.

### Cellular stiffness and fluidity as a function of temperature

The ability to measure cellular properties at constant or varying temperatures is a significant advantage of the AFS. The relatively small internal volume translates into rapid temperature equilibration. Here, we show that HEK293T cells alter their behavior in a temperature dependent manner. Cells initially removed from the incubator display surface activity; filopodia extension, fluctuation and interaction with silica particles. When cooled to 21 °C, all activity ceases, filopodia is retracted and the beads remain stationery (**Figure 4a**). We use the AFS to probe the viscoelastic properties of HEK293T cells at two different temperatures and observe marked changes in cell stiffness and fluidity. At 21 °C, HEK293T cells are significantly more compliant (E_0_ = 5.73⋇1.04 Pa) than cells measured at 37 °C (E_0_ = 9.75⋇0.8 Pa) at an applied force clamp of 58 pN (**Figure 4b**). Changes in cellular elasticity as a function of temperature is a well-documented phenomenon, observed in a variety of cells. Interestingly, how the cells respond to the change in temperature appears to be cell and/or technique dependent, with reports of decreasing stiffness with decreasing temperature^36^, or the inverse, a decrease in stiffness with increasing temperature^37–42^. These differences arise from a number of sources including measurement technique, cell type and surface coating, just to name a few. It has been speculated that these changes may be partly due to how the cell is probed, the simple act of binding a bead to a cell has been shown to induce localized actin-accumulation^43^, thereby impacting stiffness measurements.

**Figure 4.**
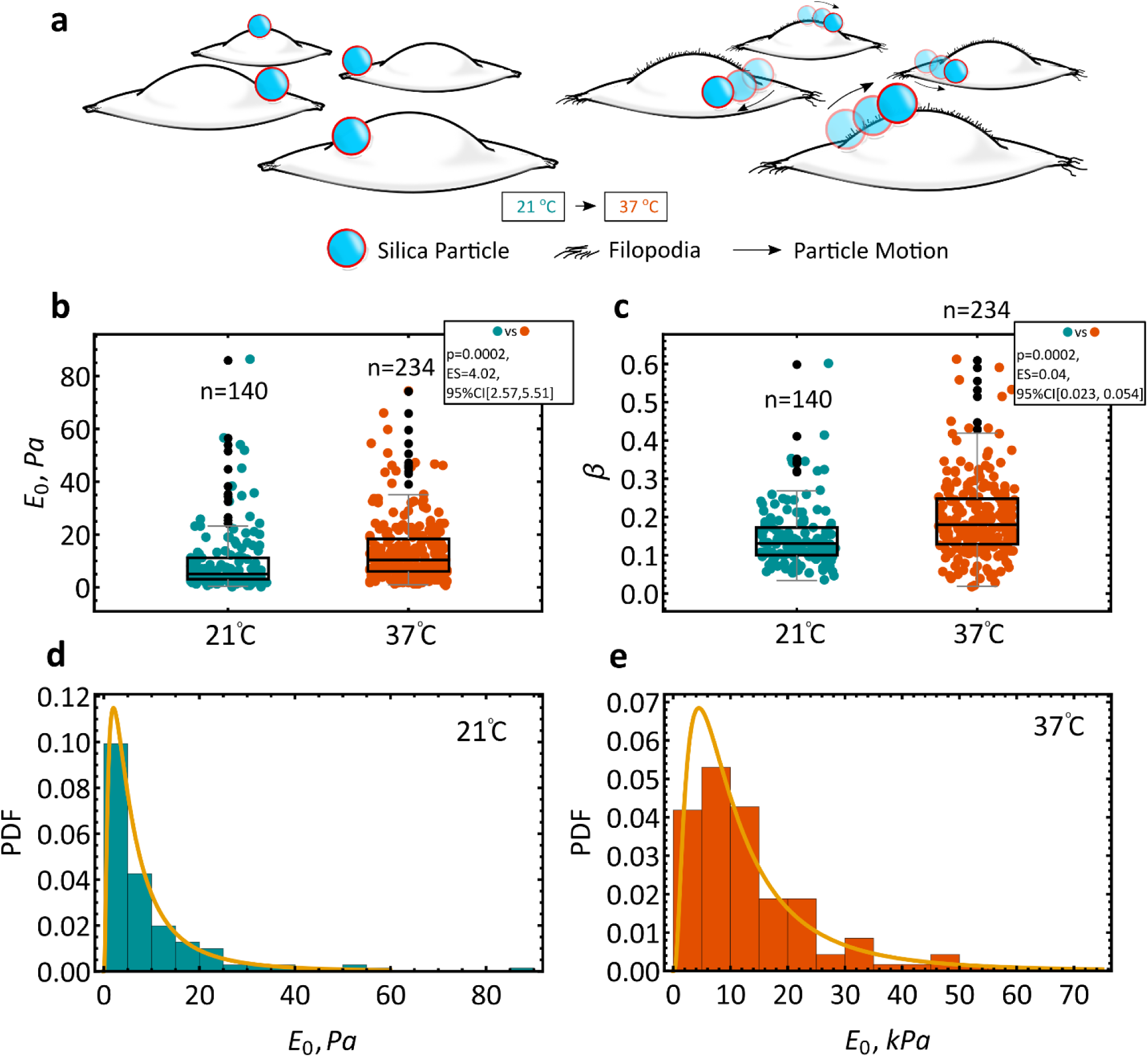
Temperature dependence. **a)**, Cells initially removed from the incubator are actively interacting with their environment. As the cells cool to 21 °C all activity ceases. Once brought back to a physiologically relevant temperature (37 °C), all activity resumes (e.g. filopodia extension and bead displacement). **b)**, The modulus scaling parameter, E_0_, is plotted as function of temperature. Cells are first measured at RT and are then brought up to 37 °C before subsequent testing. Median values of E_0_ show statistically significant differences between cells measured at 21 °C (E_0_ = 5.73⋇1.04 Pa, n = 140) and cells measured at 37 °C (E_0_ = 9.75⋇0.8 Pa, n = 234). **c)**, A statistically significant difference between the power-law exponent, β, is found for cells at 21 °C (β = 0.13⋇0.0064, n = 140) and cells at 37 °C (β = 0.17⋇0.0065, n = 234). **d)** and **e)**, The elastic modulus is best described by a log-normal distribution for all cell lines explored in this study. A force clamp of 58 pN (10% Amp) is used for all experiments. Values are reported as the geometric mean ⋇ S.E. Black dots indicate outliers. Statistics are displayed in each figure and list the p-value, effect size (ES) and the corresponding 95% confidence interval. Statistical differences are determined using estimation statistics (see Methods). Unless specified, all experiments are performed at 37 °C.

In addition, we observe a statistically significant difference between β for cells measured at 21 °C (β = 0.13⋇0.0064) and cells measured at 37 °C (β = 0.17⋇0.0065) (**Figure 4c**). Further, the distribution of the elastic modulus has been shown to vary across a single cell and is best described by a log-normal distribution^18^. The data shown in **Figure 4d** & **4e** captures the distribution of the elastic modulus across many different cells, measured at different positions. With a large enough dataset, the same log-normal distribution eventually emerges.

Taken together, our results show that as the HEK293T cell cools its stiffness reduces (lower E_0_) while at the same time exhibiting more solid-like features (decrease in β). When the microfluidic chamber is heated back to physiological temperatures the cell exerts more pull-back on the acoustically driven bead, leading to a measured increase in stiffness while at the same time exhibiting more fluid-like properties. The results obtained here are in line with those obtained by optical pulling^44^. However, the response observed here is different when compared to techniques based on indentation. AFM studies have shown that when the stiffness increases, the fluidity of the cell decreases^33,45^, that is, by becoming more solid-like, the cell increases its stiffness. While most of these experiments are performed at room temperature, some studies have looked at the stiffness and fluidity coupling at physiological temperatures. Magnetic Twisting Cytometry (MCT) on human airway smooth muscle cells showed that as the temperature increases, cells become softer (decreasing E_0_) while also becoming more fluid-like (increasing β)^40^. Probing human alveolar epithelial cells by AFM demonstrated that as the temperature increases, cells become stiffer, and also more solid-like, correlating with a reduction in viscosity^36^. Alternatively, our results corroborate early findings on immortalized murine CH27 lymphoma cells^44^ measured in suspension; as the temperature increases, the cell undergoes fluidization leading to an increase in stiffness.

The behavior observed here may arise from the fundamental difference in force directionality, that is to say, differences between pushing versus pulling. It has previously been speculated that force directionality significantly alters the mechanical effects being captured^46^. While pulling rather than pushing has been demonstrated to sensitize mechanosensitive channels^47^. The probability of pulling a tether from the plasma membrane depends on several factors, including, membrane rigidity and tension, membrane adhesion and membrane viscous resistance^48^. The coupling of the plasma membrane to the cytoskeleton leads to additional energetic considerations including the energy required to separate the cell membrane from the cytoskeleton, the energy required to bend the membrane as it is pulled into a tether with constant curvature and the energy required to overcome the viscous resistance within the plasma membrane and between the inner monolayer and the cytoskeleton^49^. As such, the response observed here is in part due to the larger contributions of these factors during pulling as opposed to indentation. For instance, the highest force used in this study of 110 pN applied over a period of 8 sec does not lead to membrane separation from the cytoskeleton. For HEK293 cells this separation has been shown to occur at a tether length of about 1.8 µm^30^. Here, at 110 pN the maximum tether length is 0.85 ± 0.10 µm demonstrating that the plasma membrane is still attached to the underlaying cytoskeletal structure. If separation does occur, then the AFS can track it (**Supplementary Figure S4**). Studies looking at fluidization and its impact on stiffness at physiological temperatures, are limited. To probe if the effects observed here are technique dependent and to draw further comparisons with published reports, we also probed the viscoelastic properties of HEK293T cells exposed to different pharmacological reagents.

#### Effect of cytoskeletal ablation on the viscoelastic properties of HEK293T cells

The cytoskeleton plays an integral part in force transmission, cellular structural stability, internal tension generation and plays a critical role in adhesion, migration and division processes^50^. These processes are driven by many factors, one of them being the cortical cytoskeleton, a filamentous actin meshwork, in contact with the plasma membrane. The polymerization of actin against the plasma membrane is involved in the generation of plasma membrane protrusions or invaginations^51^.

Here, we exposed cells to two different pharmacological treatments. First, we probe the effect of cytochalasin D on the viscoelastic properties of HEK293T cells. Cytochalasin D catalyzes the depolymerization of actin filaments^52^. A number of studies using a variety of measuring methods have demonstrated that F-actin disruption leads to an overall reduction in cell stiffness^52–56^, or inversely, cells become much more elastic and as such, easier to deform. An advantage of the AFS chip is the small microfluidic chamber which allows for rapid exchange of solutions. Cells are measured before the addition of drug followed by rapid solution exchange for the drug of choice. After 30-minute incubation with 10 µM cytochalasin D (CytoD) cells undergo a drastic change in morphology, taking on a distinctly taller, spindlier appearance (**Figure 5a**). In line with previous reports, we observe the same trend here, addition of CytoD causes a reduction in E_0_ from 11.1⋇1.29 Pa (n=171) to 7.42⋇1.25 Pa (n=191) (**Figure 5b**). Similarly, a reduction in stiffness leads to an increase in fluidity as captured by the power law exponent, β, with an increase from 0.16⋇0.0067 to 0.20⋇0.0084 (**Figure 5c**).

**Figure 5.**
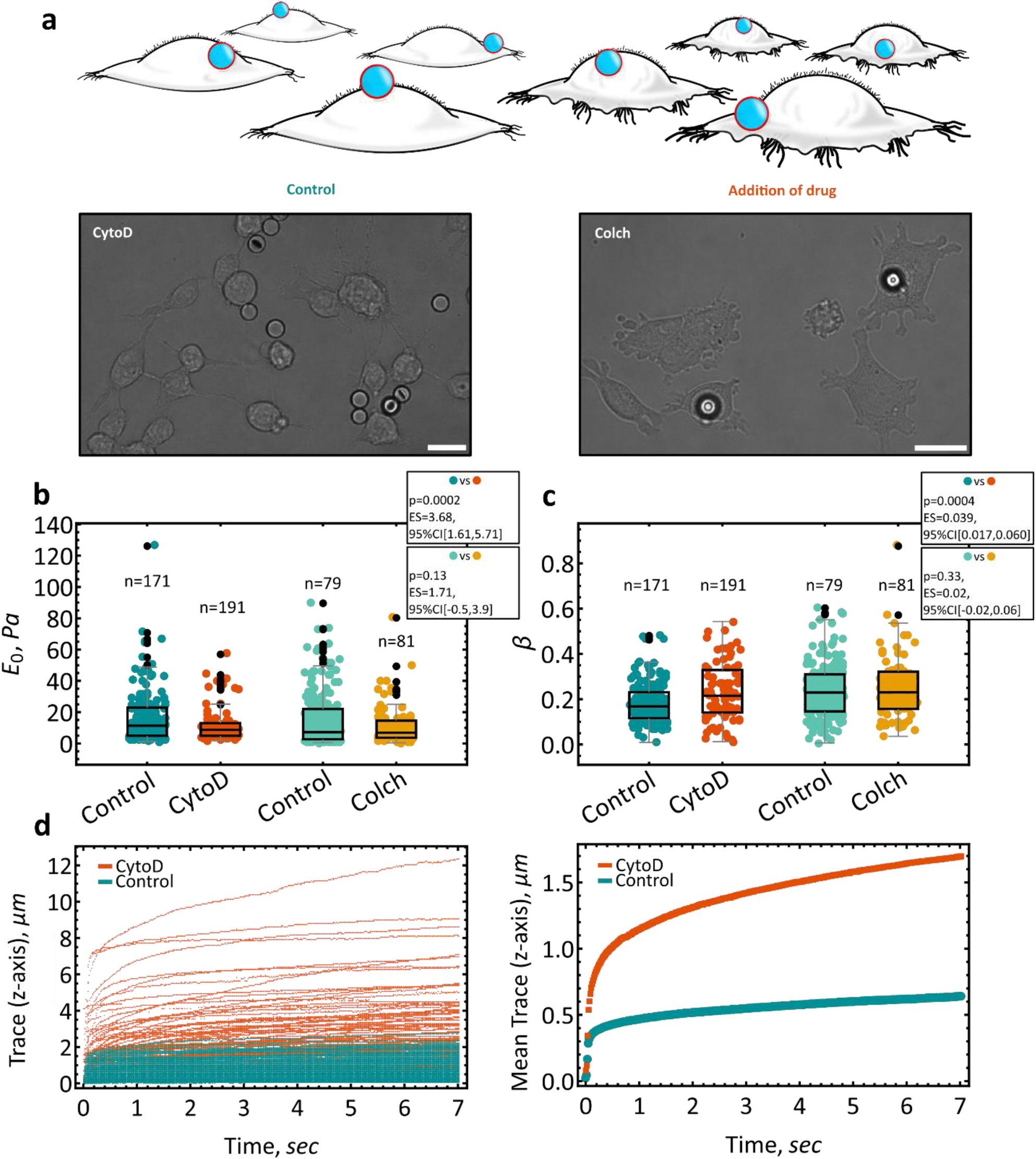
Effect of cytoskeleton modifying agents on the viscoelastic properties of HEK293T WT cells. **a)**, Cartoon of cells before addition of drugs and 30 minutes after the addition of Cytochalasin D (CytoD). Micrographs of HEKT293T cells exposed to either CytoD or Colchicine (Colch). Scale bar, 20 µm. **b)**, Addition of 10 µM CytoD leads to dramatic softening of the membrane within a span of 30 minutes (before, E_0_ = 11.1⋇1.29 Pa, n = 171), (after, E_0_ = 7.42⋇1.25 Pa, n = 191). No statistically significant differences in stiffness is found after the addition of 5 µM Colchicine (before, E_0_ = 8.84⋇1.25 Pa, n = 79), (after, E_0_ = 7.12⋇1.44 Pa, n = 81). **c)**, Statistically significant differences in the power-law exponent β are found after the addition of CytoD (before, β = 0.16⋇0.0067, n = 171), (after, β = 0.20⋇0.0084, n = 191). While, no statistically significant differences are found in the fluidity after the addition of Colchicine (before, β = 0.189⋇0.014, n = 79), (after, β = 0.209⋇0.015, n = 81). **d)**, Extension curves: control cells and after the addition of CyotD. Mean traces of both datasets show a clear difference in elasticity to the same force. A force clamp of 58 pN (10% Amp) is used for all experiments. Values are reported as the geometric mean ⋇ S.E. Black dots indicate outliers. Statistics are displayed in each figure and list the p-value, effect size (ES) and the corresponding 95% confidence interval. Statistical differences are determined using estimation statistics (see methods). Unless specified, all experiments are performed at 37 °C.

After breaking apart the smaller filamentous structures we test the impact of colchicine on microtubules (24 nm, hollow tubes). Colchicine has been shown to disrupt microtubules within neutrophils^57^ and porcine aortic endothelial cells^54^ however it appears to not affect the overall viscoelastic properties of human articular chondrocytes^52^, cancer cells (i.e. MCF-7)^58^ or fibroblasts^59^. The effect of microtubule disassembly appears to be cell dependent. Since microtubules are distributed throughout the cells stronger indentation via AFM may be required in order to characterize the effect of the drug by probing deeper into the cell^36^. By pulling on the cell surface the AFS can interrogate the surface, but not the deeper underlying structure. As such, we find no statistically significant differences between the elasticity of cells before or after treatment with 10 µM Colchicine, (E_0, before_ = 8.84⋇1.25 Pa, n = 79) and E_0, after_ 7.12⋇1.44 Pa (n = 81) (**Figure 5b**). Likewise, no differences are observed in the power law exponent, β_before_ = 0.189⋇0.014 and β_after_ = 0.209⋇0.015 (**Figure 5c**).

The drug-induced changes demonstrate the potential of the AFS system to measure the viscoelastic properties of cells in a high-throughput fashion. The impact of cytochalasin D and colchicine on fluidity and subsequently stiffness, follow previously published reports. As such, the AFS is capable of measuring the interaction of the plasma membrane with the underlying cytoskeleton and monitor perturbations to the system as is evident in **Figure 5d**. All extension traces for the control and CytoD experiments are plotted, demonstrating the significant change in overall elasticity when the underlaying actin-cortex is decoupled from the plasma membrane.

#### Effect of Piezo1 overexpression on the viscoelastic properties of HEK239T cells

Next, we employed the AFS to measure E_0_ and β of cells overexpressing a transmembrane, mechanosensitive protein, specifically, Piezo1 (**Figure 6a**). First, we re-introduced Piezo1 into a Piezo1 Knockout (KO) line using antibiotic selection and a human Piezo1-1591-GFP construct. We confirmed the overexpression of Piezo1 via the patch-clamp technique and compared the mRNA levels using quantitative real-time PCR (qPCR). The Piezo1 overexpressing line had about 70 times more Piezo1 mRNA than the WT HEK293T cells (**Figure 6b**). We employed cell-attached patch clamping to demonstrate that the Piezo1 channels are functional in the overexpressed cells (**Figure 6c and 6d**). We are able to make direct comparisons between the over-expressor (OE) and KO cell lines since they have the same origin. We include comparisons here between all 3 cell lines for completeness.

**Figure 6.**
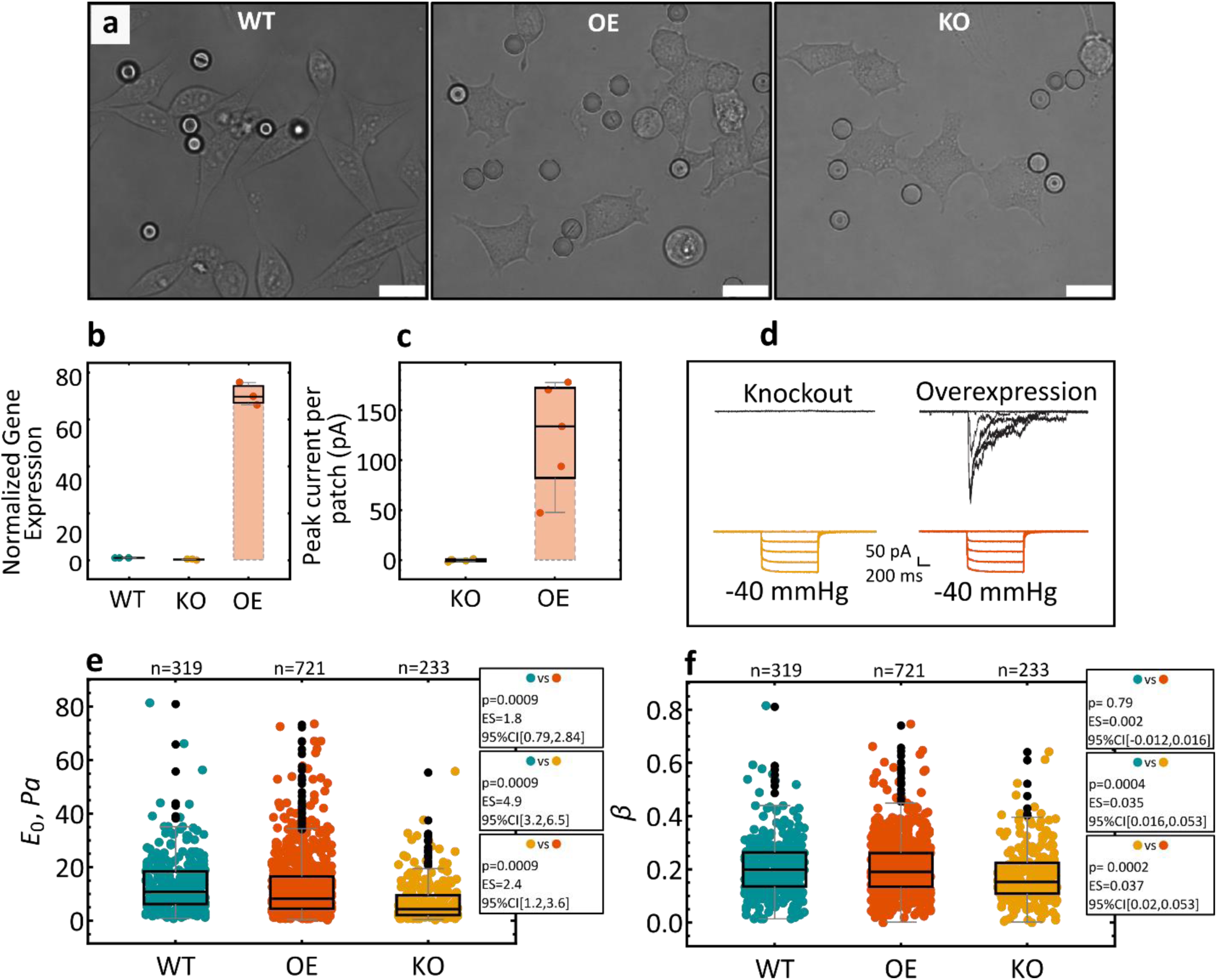
Effect of Piezo1 overexpression on the viscoelastic properties of HEK239T cells. **a)**, Micrographs showing wildtype (WT) HEK293T overexpressing Piezo1 (OE). Scale bar, 20 µm. **b)**, Normalized gene expression of Piezo1 cDNA (N=3). **c)** and **d)**, Cell-attached patch-clamping of knockout and Piezo1 overexpressing cells. **e)**, Stretching of the cell membrane by an acoustically driven, fibronectin-coated bead revealed statically significant differences in the modulus scaling parameter, E_0_, between all three cell-lines, WT (E_0_ = 10.1⋇0.57Pa, n = 319), Piezo1 Overexpressing cells (E_0_ = 8.28⋇0.40 Pa, n = 721) cells and Piezo1 knockout cells (E_0_ = 4.49⋇0.50 Pa, n = 233). **f)**, Power-law exponent of WT (β = 0.18⋇0.006, n = 319), Piezo1 Over-Expressing (β = 0.18⋇0.004, n = 721) and Piezo1 knockout cell lines (β = 0.14⋇0.007, n = 233). A force clamp of 58 pN (10% Amp) is used for all experiments. Values are reported as the geometric mean ⋇ S.E. Black dots indicate outliers. Statistics are displayed in each figure and list the p-value, effect size (ES) and the corresponding 95% confidence interval. Statistical differences are determined using estimation statistics (see methods). Unless specified, all experiments are performed at 37 °C.

We probe the mechanical properties of the three cell lines and find statistically significant differences in stiffness across all lines (**Figure 6e**). The Piezo1 knockout cells are about 80% softer than the line overexpressing Piezo1 and exhibit a more elastic-solid like behavior, as measured by a 30% decrease in fluidity (**Figure 6f**). As such, overexpression of Piezo1 leads to an increase in both the apparent stiffness and the fluidity of the cells. The measure of fluidity has been proposed as inversely related to intracellular friction which arises from the underlaying polymer network characterized by the cytoskeleton and the actomyosin complex^60^. Some have proposed that it represents the turn-over dynamics of cytoskeletal proteins^18^ where the fluidity that arises from the cytoplasm can be compared to that of a soft-glassy material (SGM). SGMs are viscous complex mixtures (emulsions, pastes and foams) that are inherently soft, disordered and metastable^45^. We extend the utility and universality of this observation to show that it can also be used to capture membrane dynamics based on protein expression levels.

While Piezo ion channels are a recent addition to the family of mechanosensitive proteins, these ion channels have already been closely linked to many mechanotransduction processes in mammalian cells^61^. For example, global knockout of Piezo1 in mice has been shown to lead to embryonic lethality^62^. Much is still unknown about how the protein activates and its various structural conformations^63,64^. Nevertheless, a recent study has shown that *Drosophila* Piezo acts to regulate mitosis while also leading to tissue stiffening as measured by AFM^65^. *Drosophila* Piezo knockout lines were shown to be softer than the same line expressing wildtype human PIEZO1. Our results demonstrate a similar trend, where the addition of Piezo1 leads to an increase in stiffness. The mechanism behind the observed stiffening will be the focus of a future study.

## CONCLUSION

Here we extend the applicability of the AFS by demonstrating high-throughput measurements of the viscoelastic properties of cells. Our results capture the heterogeneity inherent to all cells by measuring the stiffness and fluidity of cells exposed to thermal changes and pharmacological treatments. The AFS presents as a powerful tool for capturing the heterogeneity of living cells through rapid, high-throughput measurements of cellular viscoelasticity. A significant advantage of the technique is the ‘plug-and-play’ format that makes force calibration and viscoelastic measurements intuitive and easy.

## EXPERIMENTAL SECTION

### Materials

Fibronectin was purchased from Sigma Aldrich (F1141-5MG). Tygon tubing was purchased from Cole-Parmer (0.020” x 0.060"OD).

### Bead coating

Silica particles were purchased from Cospheric (9.2 µm, SS05003). Particles were first diluted in Bleach at 10 % w/v and allowed to sit for 5 minutes. Particles were washed multiple times with 1xPBS (1 mL), by centrifugation (x500g for 20 sec) and discarding the supernatant. Finally, 1 mL DMEM with 10 %FBS was added to the precipitate, for a total volume of 1 mL. Before experimental use, particles were coated with Fibronectin at a final Fibronectin concentration of 10 µg/mL in DMEM with 10%FBS and placed onto a rotating stage for 60 min at room temperature.

### Cell preparation

HEK293T cells were cultured with Dulbecco’s High-Glucose Media (DMEM) supplemented with 10% Fetal Bovine Serum (FBS) inside a humidity-controlled incubator set to 37 °C with 5% CO_2_. Cells were cultured until 90 – 95% confluent. Following Trypsinization, 300 µL of cells were spun down and supernatant removed. Fibronectin coated particles were added to the cells, with the final volume brough up to 100 µL. The cell-particle mixture was loaded into a Fibronectin coated microfluidic device (AFS chip) with a total cell coverage of around 60 - 80% (i.e. high enough density to facilitate efficient measurements without causing cells to clump). The microfluidic device was then placed into the incubator (set to 37 °C with 5% CO_2_) for 3 hours.

### Quantifying Piezo1 gene expression levels

Total RNA was isolated from HEK293T cell lines using a standard Chloroform/Propanol/Ethanol precipitation protocol. For qPCR one microgram of total RNA from each cell line in triplicate was reverse transcribed with Super Script III First Strand Synthesis Super Mix Kit (Invitrogen, Carlsbad, California). The relative abundance of Piezo1 cDNA fragments was determined using a SYBR-Green Master Mix (Roche, Basel, Switzerland). Accumulation of polymerase chain reaction products and the threshold cycle were determined using a Bio Rad CFX 384 Real Time System, C1000 Touch Thermal Cycler.

### HEK293T Piezo1 overexpression

The Piezo1 KO HEK293T line was generated by CRSIPR/Cas9. Piezo1-1591-GFP was re-introduced into this line using antibiotic selection (pcDNA3.1 vector – Geneticin 0.8 mg/ml).

### Coating and loading AFS device

Clean microfluidic devices are first rinsed with 1xPBS followed by the addition of 100 µg/mL Fibronectin. The device is then left inside the hood at room temperature for 60 minutes. To reduce the risk of introducing air bubbles into the microfluidic chip, a piece of Tygon tubing was inserted into the inlet. We used this tubing to load Fibronectin and when ready to load the cells. Tubing allows for a direct visual check for the presence of bubbles.

### Determining acoustic radiation force

The force generated by the acoustic field within the chamber can be determined by a simple force balance around the bead:

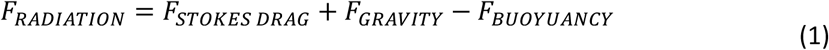

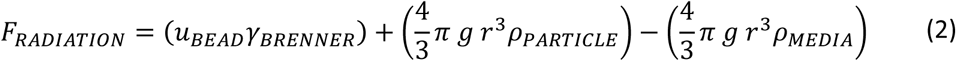

with^66^,

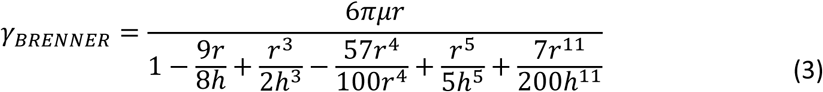

where *g* is gravity, *r* is the bead radius, *ρBEAD* is the density of the silica particle, *ρMEDIA* is the density of the cell media, *uBEAD* is the bead velocity, *γBRENNER* is the correction factor for stokes drag coefficient in close proximity to walls, *μ* is the viscosity of the media and *h* is the height of the bead center to the surface^67^. Brenner’s drag coefficient can be determined in two ways, 1) by measuring experimental drag as a function of height, where the trajectory of a particle dropped (released) from various heights is recorded and fitted using Equation 3 to obtain 6π*μr* (Stokes Drag Coefficient), used as a free-fit parameter^66^ or 2) by measuring the viscosity of the bulk fluid and directly inserting it into Equation 3.

### Determining sample viscosity

In order to find the experimental local viscosity, η, and use it for the determination of the effective drag coefficient, *γBRENNER*, we employed a method based on Brownian motion. The diffusion coefficient, *D*, is determined by measuring the Mean Square Displacement (MSD) of 1 µm resin particles as they gravitate towards the bottom of the microfluidic chamber:

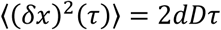

where *δx* is the x-plane movement, *d* is the dimensionality and *τ* is the time lag.

By using the Einstein relation^68^ where *T* is the temperature in Kelvin, *kB* is Boltzmann’s constant and

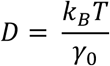

*γ*0 is bulk Stokes drag coefficient, we are able to determine the experimental viscosity using the relation *γ*0 = 6πη*r*.

### Determining the viscoelastic properties of cells

We determine the viscoelastic properties by using the power-law model, where the creep compliance of the material is captured by:

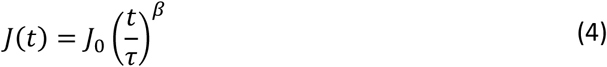

where *J*0 is the material compliance, *τ* is used normalizing time (*t*), usually set to 1 s and *β* is the power-law exponent. Values of *β*∼1 represent a more viscous material that undergoes plastic deformation, whereas values of *β*∼0 represent a more elastic material, that readily springs back to its original shape^45^. The value of β, falls in between 0.1 and 0.5 for most cells. The modulus scaling parameter, *E*0, is found by, *E*0 = 1/*J*0 and has units Pa.

The extension curves, z-height vs time, are first converted to J(t), where^24^:

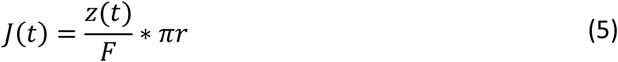

where *z*(*t*) is the extension curve (z-direction) obtained by pulling on a cell, *F* is the applied force and *r* is the particle radius. *J*(*t*) is then plotted as a function of time and Equation 4 is fit to the resultant curves. A Mathematica program was written to automatically fit the power-law to all measured distance vs time plots. Fits with an R^2^ value of lower than 0.99 are discarded. No additional filtering was used.

### Determining k_1_

The elastic coefficient is determined by fitting the distance vs times curves to the Burger’s (Spring-Dashpot) model, where:

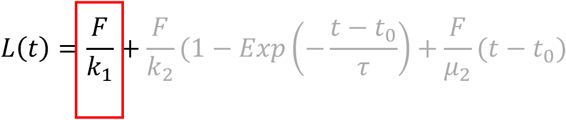

where F is the force clamp, typically 29.3 pN and *k*1 is the elastic coefficient. This component of the model describes the initial elastic response. A Mathematica code was written to extract the initial elongation distance by measuring bead displacement and location before and after the application of force. This phase finished when the bead stopped accelerating towards the node and is typically related to the maximum velocity. By determining the maximum velocity (by taking the derivative of distance vs time) and establishing an initial baseline, the initial elastic elongation (*L*0) could be extracted, 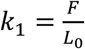.

The second part of the equation (grey) can be fit to the rest of the distance vs time plot where the particle is no longer accelerating. We do not use the Spring-Dashpot model to characterize the viscous response of the system as: “…*dissipation phenomena inside the cell cannot be assigned to a single time relaxation, nor to a finite number of time relaxations. Practically, this means that it is not possible to accurately model the cell’s mechanical behaviour by associating several springs and dashpots, respectively accounting for elastic restoring forces and viscous damping. In other words, a simple picture of the cellular medium as a Maxwell viscoelastic liquid or a Kelvin–Voigt solid, or any finite combination of them, is not realistic”* ^50^. The Spring-Dashpot model is applicable over small time ranges but has been shown to deviate over large time spans^69^.

### A note on E_0_ and β heterogeneity

Heterogeneity in both the elastic modulus and the fluidity parameter can arise from a number of sources including the cell cycle, acoustic force variation across the chip and variation in bead diameter used for determining acoustic force.

### Time dependence

Cell measurements are started close to the inlet. A single field-of-view is chosen close to the inlet, starting on one side of the channel. Cells are probed and extension curves are recorded. The field-of-view (FoV) is then manually shifted (left or right, depending on which side of the channel the measurements are started). Here, the FoV is shifted to the left. We proceed to probe the cells within the next field of view. Once done, the FoV is moved left again, by one FoV. Measurements are carried out for 30 minutes, which roughly equals about a 1/3 of the microfluidic channel length. Measurements are made for another 45 minutes (which represents the 75 min. mark) and cover the second 1/3 of the channel. After another 45 minutes of measurements (data represents the 120 min. mark), the channel outlet is typically reached.

### Temperature dependence

Cells, initially at 37 °C are taken out of the incubator, placed onto the AFS stage and are allowed to cool to RT (21C) for 15 minutes. Once cooled, extension (elongation) curves are obtained over a period of 30 minutes. Next, the cells are heated back to 37 °C. After 15 minutes, the cells are measured again for 30 minutes.

### Addition of pharmacological reagents

All reagents were diluted and used on the day of the experiment. Cytochalasin D (CytoD) was diluted in DMEM with 10% FBS to a final concentration of 10 µM. This mixture was kept at 37 °C using a water bath. The viscoelastic properties of cells were first measured for 30 minutes prior to the addition of the drug. After 30 minutes, CytoD was introduced into the microfluidic channel, completely replacing all media. (Importantly, before addition of a new reagent the temperature controller has to be turned off. If the temperature is set to 37 °C while injecting new media that is slightly lower than this set point, the temperature controller will try to compensate, resulting in cellular death.) The device was then placed into an incubator for 30 minutes and was then placed back onto the stage with the temperature controller set to 37 °C. Cells were then probed for another 30 minutes. Colchicine was diluted in DMEM with 10 %FBS to a final concentration of 5 µM. Similar to the protocol for CytoD, cells were however, incubated with the drug for about 5 – 10 minutes before probing. Several Colchicine concentrations were tested, 5 µM to 50 µM, with no apparent change in viscoelastic properties (data not shown).

### General data acquisition

Data were acquired using the LUMICKS AFS stand-alone technology including a LabVIEW interface for microsphere tracking. All measurements (unless specified) were performed within 60 minutes at 37 °C thanks to the integrated temperature control (AFS-TC, LUMICKS) with an amplitude of 10% at 14.45 MHz frequency, applied constantly for a period of 8 secs.

Images were acquired with a bright-field inverted microscope equipped with 1.3 MP camera recording at 60 Hz (UI-324CP, IDS). An air 20x 0.75 NA objective (Nikon, CFI Plan APO, VC 20x, (MRD70200)) was used for tracking bead movement.

### Patch-clamp electrophysiology

Cell-attached patch-clamp recordings were carried out at room temperature as previously described^70^. The extracellular solution for cell-attached patches contained high K^+^ to zero the membrane potential and consisted of 90 mM potassium aspartate, 50 mM KCl, 1 mM MgCl_2_ and 10 mM HEPES (pH 7.2) adjusted using KOH. The pipette solution contained 140 mM NaCl, 1 mM MgCl_2_ with 10 mM HEPES (pH 7.2) adjusted using sodium hydroxide. Negative pressure was applied to patch pipettes using a High Speed Pressure Clamp-1 (ALA Scientific Instruments) and recorded in millimeters of mercury (mmHg) using a piezoelectric pressure transducer (WPI, Sarasota, FL, USA).

### Statistical analysis and bootstrapping

The data for E_0,_ in line with previous results, is log-normally distributed. The data for β, at large sample sizes was also best described by a log-normal distribution. To obtain mean values the data was log-transformed, where finally the geometric mean and the geometric standard error is reported. Finally, the exponential is taken of the mean and the standard error and is reported within the document. Statistical significance was tested by bootstrapping the sample 5000 times and reporting the mean difference between the samples. The full protocol is outlined elsewhere^71^. The p-value is the likelihood of observing the effect size, if the null hypothesis of zero difference is true^71^. We report the bootstrapped effect size and the corresponding 95% confidence interval. Data is taken as significant if the p-value is lower than 0.005^72^ and the corresponding effect size is well bounded within a narrow 95% confidence interval (CI). The larger the span of the 95% CI, the less confidence is placed in the p-value. Box and whisker plots display the median, 25% and 75% quartiles, with whiskers at an interquartile range of 1.5 (95% confidence). Outliers are shown as black dots.

## ASSOCIATED CONTENT

Supporting information contains the protocol for determining sample viscosity; effect of numeral aperture on particle tracking; cartoon of the data collection protocol; plot of cytoskeleton-membrane detachment.

## ACKNOWLEDGMENTS

We would like to thank Jasmina Cvetkovska for generating the qPCR data.

## SUPPORTING INFORMATION

**Figure S1.**
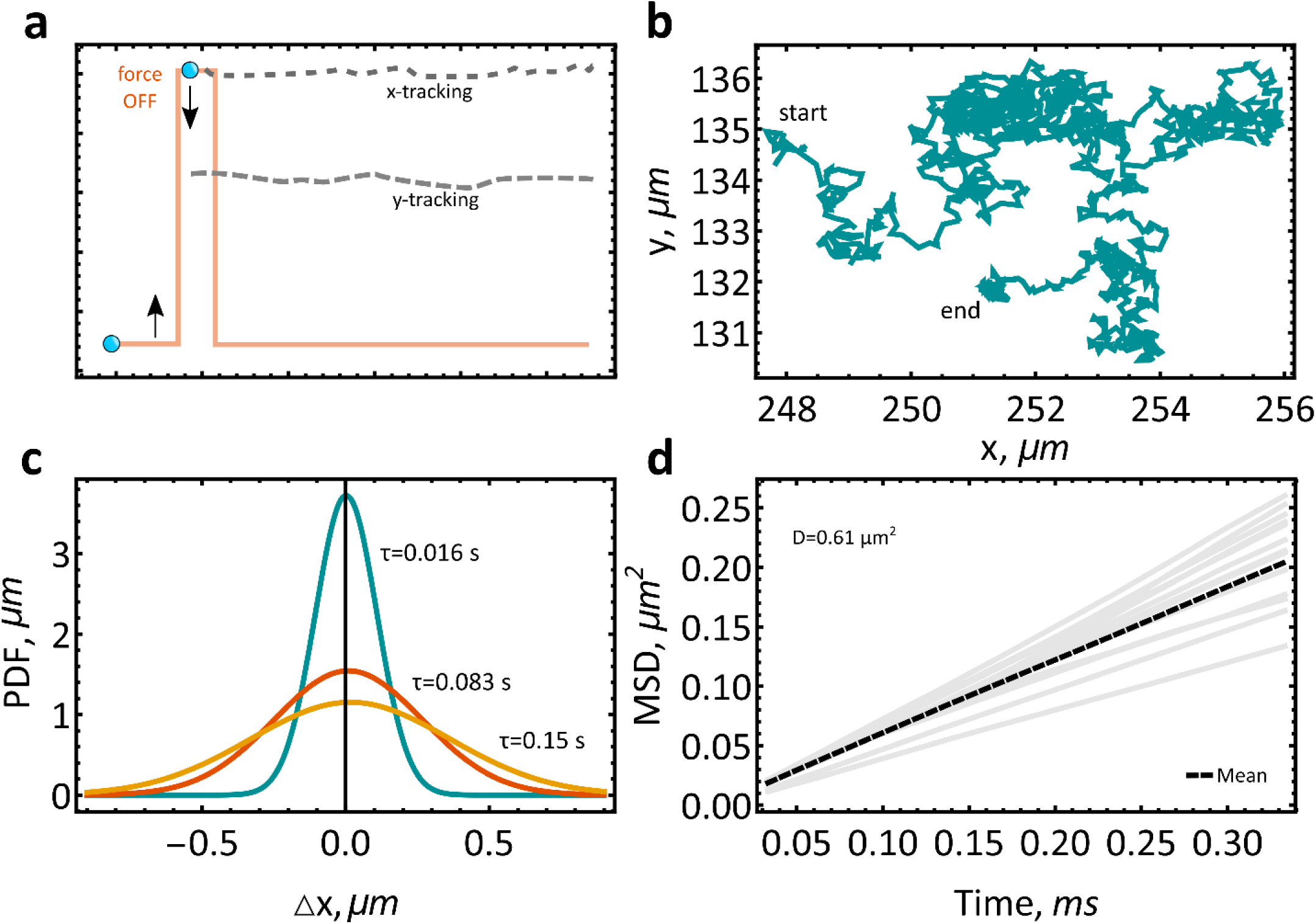
Experimental determination of viscosity. a), Silica particles (1 µm), driven to the acoustic node, undergo Brownian motion once the acoustic field is turned off. **b)**, Mean Square Displacement (MSD) is determined by tracking of the x and y component as the particles slowly settle toward the bottom of the channel. **c)**, Plotting time lag, *τ*, versus x-displacement, demonstrates that particle travel is well described by a gaussian, centered at 0 µm. **d)**, The mean diffusion, D, is calculated by taking the slope of 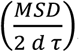, where d is the dimension ({x} = 1, {y,x} = 2) (n=27).

**Figure 2S.**
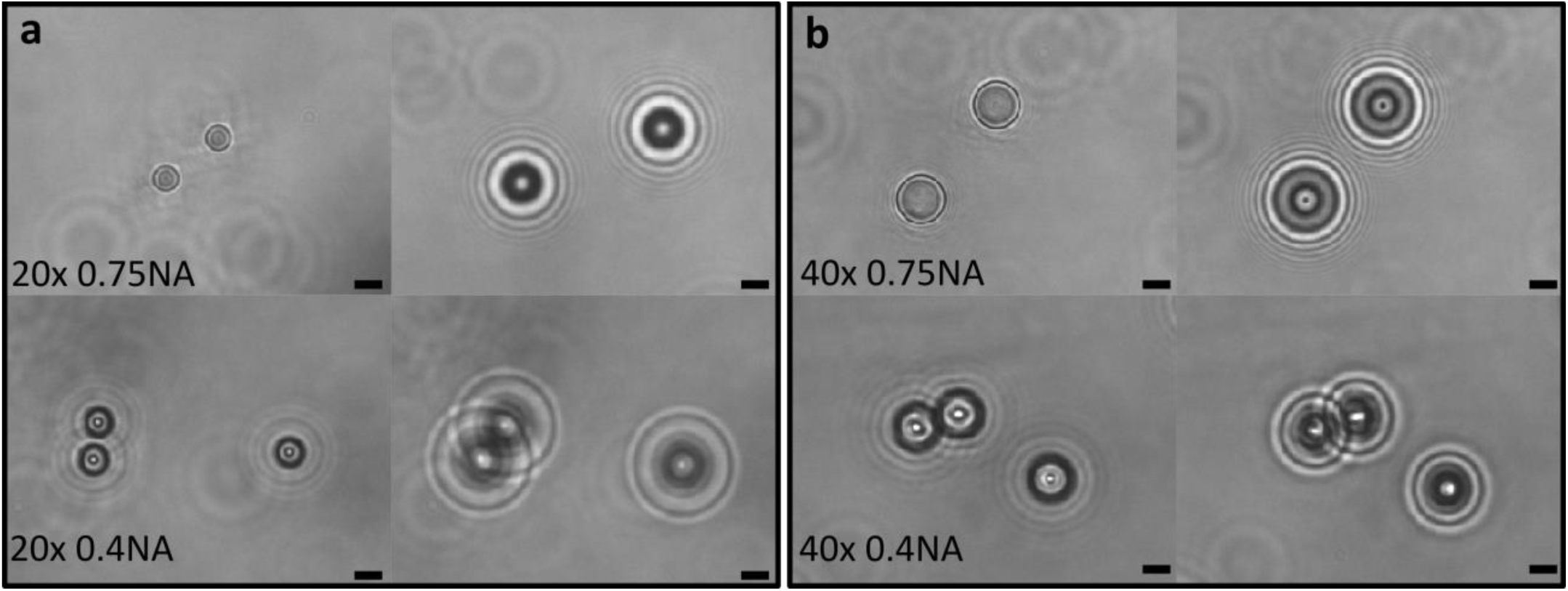
Effect of numerical aperture on particle tracking. **a)**, An objective with a high numerical aperture creates distinct airy patterns that are readily tracked by the software. A low numerical aperture objective does not create enough contrast for consistent tracking of beads when in proximity to cells. Scale bar, 10 µm. **b)** Higher magnification objectives with lower NA can be used for particle tracking at the expense of throughput. Scale bar, 5 µm.

**Figure S3.**
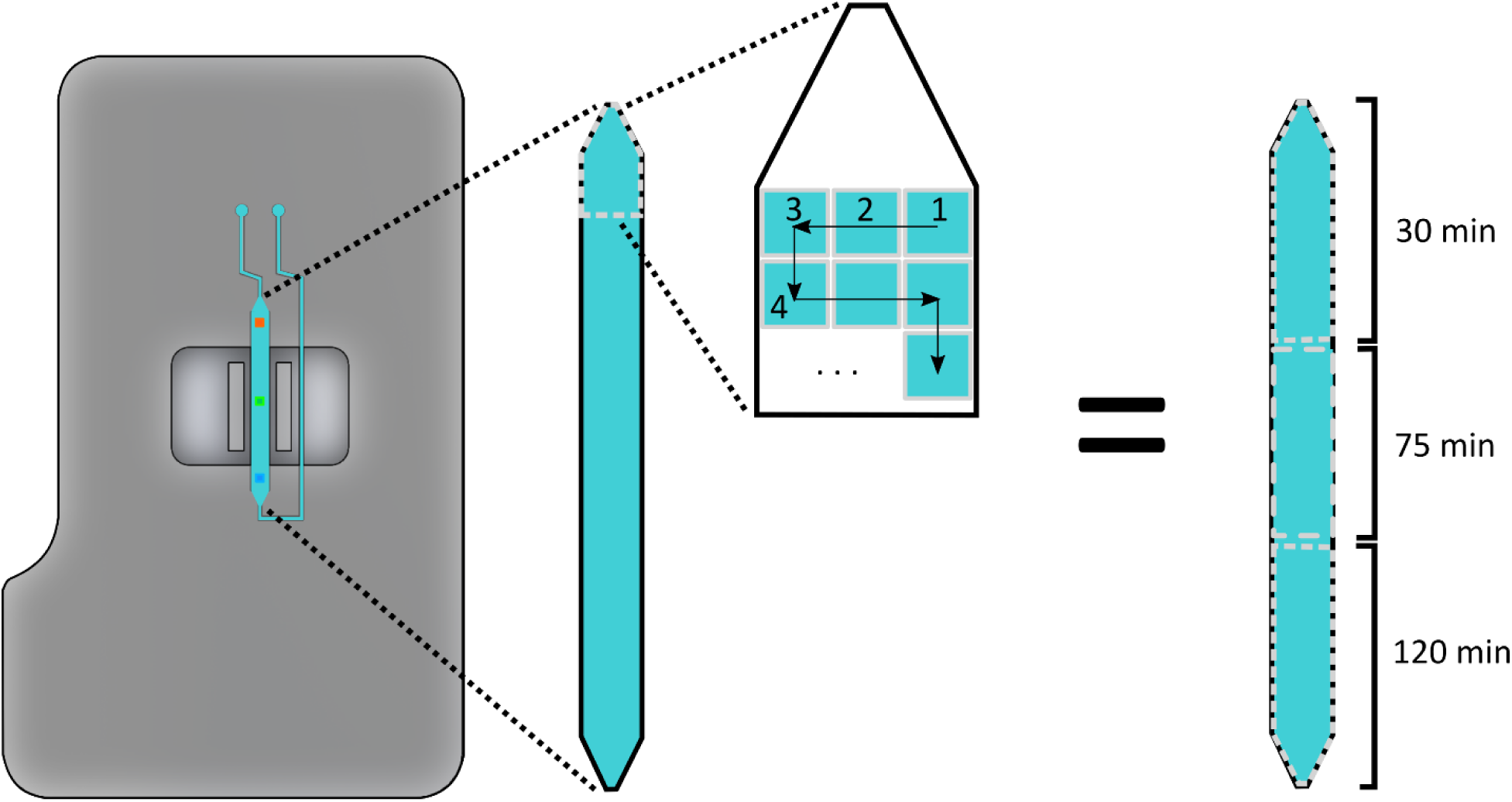
Data collection. For full protocol please refer to the Experimental section.

**Figure. S4.**
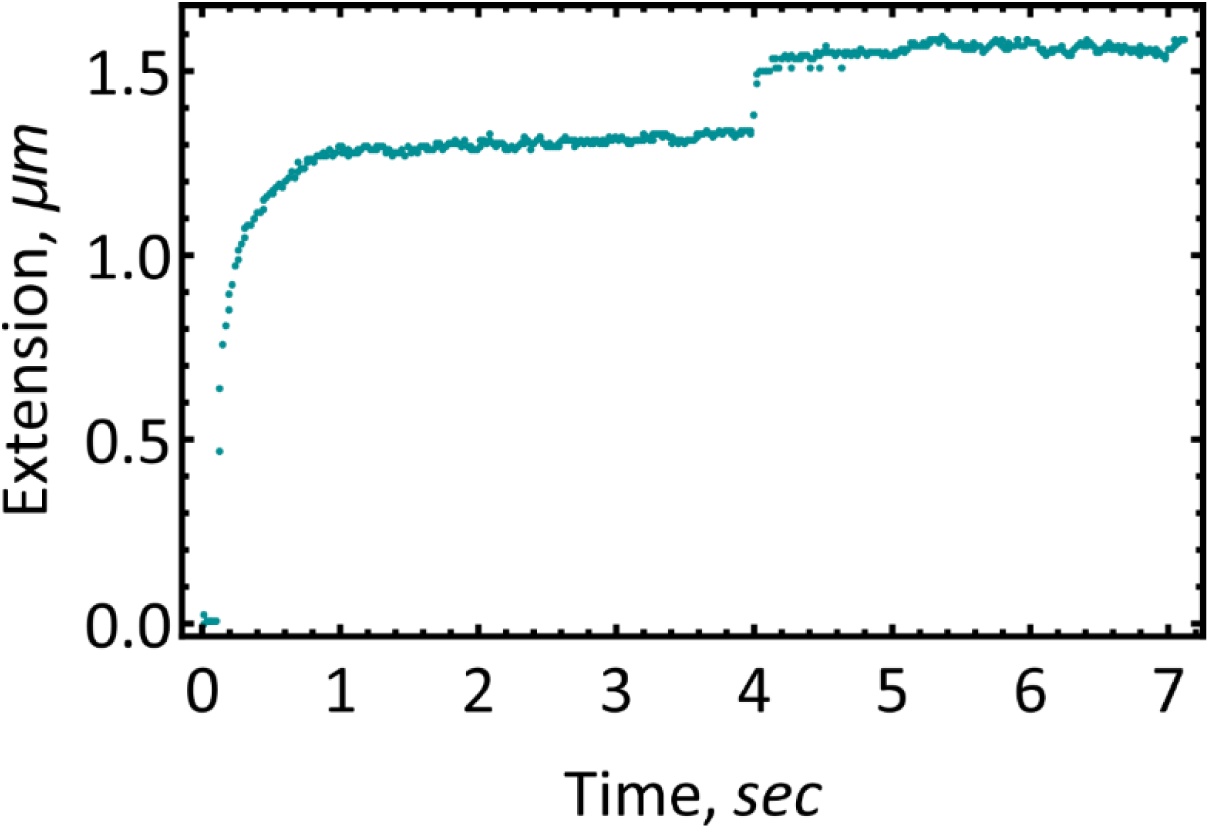
Breaking of the membrane from the underlaying cytoskeleton. The jump at around 4 sec is indicative of this separation. Force 58.5 pN (14.45MHz), HEK293 WT, 37C.

## REFERENCES

(1) Huang, H.; Kamm, R. D.; Lee, R. T. Cell Mechanics and Mechanotransduction: Pathways, Probes, and Physiology. Am. J. Physiol.-Cell Physiol. 2004, 287 (1), C1–C11. https://doi.org/10.1152/ajpcell.00559.2003.

(2) Dao, M.; Lim, C. T.; Suresh, S. Mechanics of the Human Red Blood Cell Deformed by Optical Tweezers. J. Mech. Phys. Solids 2003, 51 (11), 2259–2280. https://doi.org/10.1016/j.jmps.2003.09.019.

(3) Lee, G. Y. H.; Lim, C. T. Biomechanics Approaches to Studying Human Diseases. Trends Biotechnol. 2007, 25 (3), 111–118. https://doi.org/10.1016/j.tibtech.2007.01.005.

(4) Tse, J. M.; Cheng, G.; Tyrrell, J. A.; Wilcox-Adelman, S. A.; Boucher, Y.; Jain, R. K.; Munn, L. L. Mechanical Compression Drives Cancer Cells toward Invasive Phenotype. Proc. Natl. Acad. Sci. 2012, 109 (3), 911–916. https://doi.org/10.1073/pnas.1118910109.

(5) Krishnan, R.; Park, C. Y.; Lin, Y.-C.; Mead, J.; Jaspers, R. T.; Trepat, X.; Lenormand, G.; Tambe, D.; Smolensky, A. V.; Knoll, A. H.; Butler, J. P.; Fredberg, J. J. Reinforcement versus Fluidization in Cytoskeletal Mechanoresponsiveness. PLOS ONE 2009, 4 (5), e5486. https://doi.org/10.1371/journal.pone.0005486.

(6) Wu, P.-H.; Aroush, D. R.-B.; Asnacios, A.; Chen, W.-C.; Dokukin, M. E.; Doss, B. L.; Durand-Smet, P.; Ekpenyong, A.; Guck, J.; Guz, N. V.; Janmey, P. A.; Lee, J. S. H.; Moore, N. M.; Ott, A.; Poh, Y.-C.; Ros, R.; Sander, M.; Sokolov, I.; Staunton, J. R.; Wang, N.; Whyte, G.; Wirtz, D. A Comparison of Methods to Assess Cell Mechanical Properties. Nat. Methods 2018, 15 (7), 491–498. https://doi.org/10.1038/s41592-018-0015-1.

(7) Darling, E. M.; Di Carlo, D. High-Throughput Assessment of Cellular Mechanical Properties. Annu. Rev. Biomed. Eng. 2015, 17 (1), 35–62. https://doi.org/10.1146/annurev-bioeng-071114-040545.

(8) Ziemann, F.; Rädler, J.; Sackmann, E. Local Measurements of Viscoelastic Moduli of Entangled Actin Networks Using an Oscillating Magnetic Bead Micro-Rheometer. Biophys. J. 1994, 66 (6), 2210–2216. https://doi.org/10.1016/S0006-3495(94)81017-3.

(9) Putman, C. A.; van der Werf, K. O.; de Grooth, B. G.; van Hulst, N. F.; Greve, J. Viscoelasticity of Living Cells Allows High Resolution Imaging by Tapping Mode Atomic Force Microscopy. Biophys. J. 1994, 67 (4), 1749–1753. https://doi.org/10.1016/S0006-3495(94)80649-6.

(10) Trickey, W. R.; Lee, G. M.; Guilak, F. Viscoelastic Properties of Chondrocytes from Normal and Osteoarthritic Human Cartilage. J. Orthop. Res. 2000, 18 (6), 891–898. https://doi.org/10.1002/jor.1100180607.

(11) Mason, T. G.; Ganesan, K.; van Zanten, J. H.; Wirtz, D.; Kuo, S. C. Particle Tracking Microrheology of Complex Fluids. Phys. Rev. Lett. 1997, 79 (17), 3282–3285. https://doi.org/10.1103/PhysRevLett.79.3282.

(12) Kamsma, D.; Bochet, P.; Oswald, F.; Alblas, N.; Goyard, S.; Wuite, G. J. L.; Peterman, E. J. G.; Rose, T. Single-Cell Acoustic Force Spectroscopy: Resolving Kinetics and Strength of T Cell Adhesion to Fibronectin. Cell Rep. 2018, 24 (11), 3008–3016. https://doi.org/10.1016/j.celrep.2018.08.034.

(13) Sorkin, R.; Bergamaschi, G.; Kamsma, D.; Brand, G.; Dekel, E.; Ofir-Birin, Y.; Rudik, A.; Gironella, M.; Ritort, F.; Regev-Rudzki, N.; Roos, W. H.; Wuite, G. J. L. Probing Cellular Mechanics with Acoustic Force Spectroscopy. Mol. Biol. Cell 2018, 29 (16), 2005–2011. https://doi.org/10.1091/mbc.E18-03-0154.

(14) Nguyen, A.; Brandt, M.; Betz, T. Microchip Based Microrheology via Acoustic Force Spectroscopy Shows That Endothelial Cell Mechanics Follows a Fractional Viscoelastic Model. bioRxiv 2020, 2020.07.02.185330. https://doi.org/10.1101/2020.07.02.185330.

(15) Martinac, B.; Nikolaev, Y.A.; Silvani, G.; Bavi, N.; Romanov, V.; Nakayama, Y.; Martinac, A.D.; Rohde, P.; Bavi, O.; Cox, C.D. Cell Membrane Mechanics and Mechanosensory Transduction.; Current Topics in Membranes: Membrane Biomechanics; Elsevier, 2020.

(16) Khatibzadeh, N.; Gupta, S.; Farrell, B.; Brownell, W. E.; Anvari, B. Effects of Cholesterol on Nano-Mechanical Properties of the Living Cell Plasma Membrane. Soft Matter 2012, 8 (32), 8350–8360. https://doi.org/10.1039/C2SM25263E.

(17) Lange, J. R.; Steinwachs, J.; Kolb, T.; Lautscham, L. A.; Harder, I.; Whyte, G.; Fabry, B. Microconstriction Arrays for High-Throughput Quantitative Measurements of Cell Mechanical Properties. Biophys. J. 2015, 109 (1), 26–34. https://doi.org/10.1016/j.bpj.2015.05.029.

(18) M. Hecht, F.; Rheinlaender, J.; Schierbaum, N.; H. Goldmann, W.; Fabry, B.; E. Schäffer, T. Imaging Viscoelastic Properties of Live Cells by AFM: Power-Law Rheology on the Nanoscale. Soft Matter 2015, 11 (23), 4584–4591. https://doi.org/10.1039/C4SM02718C.

(19) Sitters, G.; Kamsma, D.; Thalhammer, G.; Ritsch-Marte, M.; Peterman, E. J. G.; Wuite, G. J. L. Acoustic Force Spectroscopy. Nat. Methods 2015, 12 (1), 47–50. https://doi.org/10.1038/nmeth.3183.

(20) Lide, D. R. CRC Handbook of Chemistry and Physics: A Ready-Reference Book of Chemical and Physical Data; CRC Press, 1995.

(21) Croughan, M. S.; Sayre, E. S.; Wang, D. I. C. Viscous Reduction of Turbulent Damage in Animal Cell Culture. Biotechnol. Bioeng. 1989, 33 (7), 862–872. https://doi.org/10.1002/bit.260330710.

(22) Wang, C.; Lu, H.; Schwartz, M. A. A Novel in Vitro Flow System for Changing Flow Direction on Endothelial Cells. J. Biomech. 2012, 45 (7), 1212–1218. https://doi.org/10.1016/j.jbiomech.2012.01.045.

(23) Wu, P.-H.; Aroush, D. R.-B.; Asnacios, A.; Chen, W.-C.; Dokukin, M. E.; Doss, B. L.; Durand, P.; Ekpenyong, A.; Guck, J.; Guz, N. V.; Janmey, P. A.; Lee, J. S. H.; Moore, N. M.; Ott, A.; Poh, Y.-C.; Ros, R.; Sander, M.; Sokolov, I.; Staunton, J. R.; Wang, N.; Whyte, G.; Wirtz, D. Comparative Study of Cell Mechanics Methods. Nat. Methods 2018, 15 (7), 491–498. https://doi.org/10.1038/s41592-018-0015-1.

(24) Kollmannsberger, P.; Mierke, C. T.; Fabry, B. Nonlinear Viscoelasticity of Adherent Cells Is Controlled by Cytoskeletal Tension. Soft Matter 2011, 7 (7), 3127–3132. https://doi.org/10.1039/C0SM00833H.

(25) Nawaz, S.; Sánchez, P.; Bodensiek, K.; Li, S.; Simons, M.; Schaap, I. A. T. Cell Visco-Elasticity Measured with AFM and Optical Trapping at Sub-Micrometer Deformations. PLoS ONE 2012, 7 (9). https://doi.org/10.1371/journal.pone.0045297.

(26) Ndoye, F.; Yousafzai, M. S.; Coceano, G.; Bonin, S.; Scoles, G.; Ka, O.; Niemela, J.; Cojoc, D. The Influence of Lateral Forces on the Cell Stiffness Measurement by Optical Tweezers Vertical Indentation. Int. J. Optomechatronics 2016, 10 (1), 53–62. https://doi.org/10.1080/15599612.2016.1149896.

(27) Yousafzai, M. S.; Ndoye, F.; Coceano, G.; Niemela, J.; Bonin, S.; Scoles, G.; Cojoc, D. Substrate-Dependent Cell Elasticity Measured by Optical Tweezers Indentation. Opt. Lasers Eng. 2016, 76, 27–33. https://doi.org/10.1016/j.optlaseng.2015.02.008.

(28) Haghparast, S. M. A.; Kihara, T.; Miyake, J. Distinct Mechanical Behavior of HEK293 Cells in Adherent and Suspended States. PeerJ 2015, 3, e1131. https://doi.org/10.7717/peerj.1131.

(29) Li, M.; Liu, L.; Xiao, X.; Xi, N.; Wang, Y. Effects of Methotrexate on the Viscoelastic Properties of Single Cells Probed by Atomic Force Microscopy. J. Biol. Phys. 2016, 42 (4), 551–569. https://doi.org/10.1007/s10867-016-9423-6.

(30) Khatibzadeh, N.; Spector, A. A.; Brownell, W. E.; Anvari, B. Effects of Plasma Membrane Cholesterol Level and Cytoskeleton F-Actin on Cell Protrusion Mechanics. PLoS ONE 2013, 8 (2). https://doi.org/10.1371/journal.pone.0057147.

(31) Ermilov, S. A.; Murdock, D. R.; Qian, F.; Brownell, W. E.; Anvari, B. Studies of Plasma Membrane Mechanics and Plasma Membrane–Cytoskeleton Interactions Using Optical Tweezers and Fluorescence Imaging. J. Biomech. 2007, 40 (2), 476–480. https://doi.org/10.1016/j.jbiomech.2005.12.006.

(32) Acerbi, I.; Luque, T.; Giménez, A.; Puig, M.; Reguart, N.; Farré, R.; Navajas, D.; Alcaraz, J. Integrin-Specific Mechanoresponses to Compression and Extension Probed by Cylindrical Flat-Ended AFM Tips in Lung Cells. PloS One 2012, 7 (2), e32261. https://doi.org/10.1371/journal.pone.0032261.

(33) Fabry, B.; Maksym, G. N.; Butler, J. P.; Glogauer, M.; Navajas, D.; Fredberg, J. J. Scaling the Microrheology of Living Cells. Phys. Rev. Lett. 2001, 87 (14), 148102. https://doi.org/10.1103/PhysRevLett.87.148102.

(34) Massiera, G.; Van Citters, K. M.; Biancaniello, P. L.; Crocker, J. C. Mechanics of Single Cells: Rheology, Time Dependence, and Fluctuations. Biophys. J. 2007, 93 (10), 3703–3713. https://doi.org/10.1529/biophysj.107.111641.

(35) Neuman, K. C.; Nagy, A. Single-Molecule Force Spectroscopy: Optical Tweezers, Magnetic Tweezers and Atomic Force Microscopy. Nat. Methods 2008, 5 (6), 491–505. https://doi.org/10.1038/nmeth.1218.

(36) Sunyer, R.; Trepat, X.; Fredberg, J. J.; Farré, R.; Navajas, D. The Temperature Dependence of Cell Mechanics Measured by Atomic Force Microscopy. Phys. Biol. 2009, 6 (2), 025009. https://doi.org/10.1088/1478-3975/6/2/025009.

(37) Sunnerberg, J. P.; Moore, P.; Spedden, E.; Kaplan, D. L.; Staii, C. Variations of Elastic Modulus and Cell Volume with Temperature for Cortical Neurons. Langmuir 2019, 35 (33), 10965–10976. https://doi.org/10.1021/acs.langmuir.9b01651.

(38) Chan, C. J.; Whyte, G.; Boyde, L.; Salbreux, G.; Guck, J. Impact of Heating on Passive and Active Biomechanics of Suspended Cells. Interface Focus 2014, 4 (2), 20130069. https://doi.org/10.1098/rsfs.2013.0069.

(39) Spedden, E.; Kaplan, D. L.; Staii, C. Temperature Response of the Neuronal Cytoskeleton Mapped via Atomic Force and Fluorescence Microscopy. Phys. Biol. 2013, 10 (5), 056002. https://doi.org/10.1088/1478-3975/10/5/056002.

(40) Bursac, P.; Lenormand, G.; Fabry, B.; Oliver, M.; Weitz, D. A.; Viasnoff, V.; Butler, J. P.; Fredberg, J. J. Cytoskeletal Remodelling and Slow Dynamics in the Living Cell. Nat. Mater. 2005, 4 (7), 557–561. https://doi.org/10.1038/nmat1404.

(41) Vlahakis, N. E.; Schroeder, M. A.; Pagano, R. E.; Hubmayr, R. D. Role of Deformation-Induced Lipid Trafficking in the Prevention of Plasma Membrane Stress Failure. Am. J. Respir. Crit. Care Med. 2002, 166 (9), 1282–1289. https://doi.org/10.1164/rccm.200203-207OC.

(42) Stroetz, R. W.; Vlahakis, N. E.; Walters, B. J.; Schroeder, M. A.; Hubmayr, R. D. Validation of a New Live Cell Strain System: Characterization of Plasma Membrane Stress Failure. J. Appl. Physiol. 2001, 90 (6), 2361–2370. https://doi.org/10.1152/jappl.2001.90.6.2361.

(43) Deng, L.; Fairbank, N. J.; Cole, D. J.; Fredberg, J. J.; Maksym, G. N. Airway Smooth Muscle Tone Modulates Mechanically Induced Cytoskeletal Stiffening and Remodeling. J. Appl. Physiol. 2005, 99 (2), 634–641. https://doi.org/10.1152/japplphysiol.00025.2005.

(44) Maloney, J. M.; Lehnhardt, E.; Long, A. F.; Van Vliet, K. J. Mechanical Fluidity of Fully Suspended Biological Cells. Biophys. J. 2013, 105 (8), 1767–1777. https://doi.org/10.1016/j.bpj.2013.08.040.

(45) Kollmannsberger, P.; Fabry, B. Linear and Nonlinear Rheology of Living Cells. Annu. Rev. Mater. Res. 2011, 41 (1), 75–97. https://doi.org/10.1146/annurev-matsci-062910-100351.

(46) Smith, B. A.; Tolloczko, B.; Martin, J. G.; Grütter, P. Probing the Viscoelastic Behavior of Cultured Airway Smooth Muscle Cells with Atomic Force Microscopy: Stiffening Induced by Contractile Agonist. Biophys. J. 2005, 88 (4), 2994–3007. https://doi.org/10.1529/biophysj.104.046649.

(47) Gaub, B. M.; Müller, D. J. Mechanical Stimulation of Piezo1 Receptors Depends on Extracellular Matrix Proteins and Directionality of Force. Nano Lett. 2017, 17 (3), 2064–2072. https://doi.org/10.1021/acs.nanolett.7b00177.

(48) Nussenzveig, H. M. Cell Membrane Biophysics with Optical Tweezers. Eur. Biophys. J. 2018, 47 (5), 499–514. https://doi.org/10.1007/s00249-017-1268-9.

(49) Li, Z.; Anvari, B.; Takashima, M.; Brecht, P.; Torres, J. H.; Brownell, W. E. Membrane Tether Formation from Outer Hair Cells with Optical Tweezers. Biophys. J. 2002, 82 (3), 1386–1395. https://doi.org/10.1016/S0006-3495(02)75493-3.

(50) Balland, M.; Richert, A.; Gallet, F. The Dissipative Contribution of Myosin II in the Cytoskeleton Dynamics of Myoblasts. Eur. Biophys. J. 2005, 34 (3), 255–261. https://doi.org/10.1007/s00249-004-0447-7.

(51) Saarikangas, J.; Zhao, H.; Lappalainen, P. Regulation of the Actin Cytoskeleton-Plasma Membrane Interplay by Phosphoinositides. Physiol. Rev. 2010, 90 (1), 259–289. https://doi.org/10.1152/physrev.00036.2009.

(52) Trickey, W. R.; Vail, T. P.; Guilak, F. The Role of the Cytoskeleton in the Viscoelastic Properties of Human Articular Chondrocytes. J. Orthop. Res. 2004, 22 (1), 131–139. https://doi.org/10.1016/S0736-0266(03)0150-5.

(53) Reynolds, N. H.; Ronan, W.; Dowling, E. P.; Owens, P.; McMeeking, R. M.; McGarry, J. P. On the Role of the Actin Cytoskeleton and Nucleus in the Biomechanical Response of Spread Cells. Biomaterials 2014, 35 (13), 4015–4025. https://doi.org/10.1016/j.biomaterials.2014.01.056.

(54) Sato, M.; Theret, D. P.; Wheeler, L. T.; Ohshima, N.; Nerem, R. M. Application of the Micropipette Technique to the Measurement of Cultured Porcine Aortic Endothelial Cell Viscoelastic Properties. J. Biomech. Eng. 1990, 112 (3), 263–268. https://doi.org/10.1115/1.2891183.

(55) Tan, S. C.; Pan, W. X.; Ma, G.; Cai, N.; Leong, K. W.; Liao, K. Viscoelastic Behaviour of Human Mesenchymal Stem Cells. BMC Cell Biol. 2008, 9 (1), 40. https://doi.org/10.1186/1471-2121-9-40.

(56) Nijenhuis, N.; Zhao, X.; Carisey, A.; Ballestrem, C.; Derby, B. Combining AFM and Acoustic Probes to Reveal Changes in the Elastic Stiffness Tensor of Living Cells. Biophys. J. 2014, 107 (7), 1502–1512. https://doi.org/10.1016/j.bpj.2014.07.073.

(57) Chien, S.; Sung, K. L. Effect of Colchicine on Viscoelastic Properties of Neutrophils. Biophys. J. 1984, 46 (3), 383–386. https://doi.org/10.1016/S0006-3495(84)84034-5.

(58) Wu, Y.; Cheng, T.; Chen, Q.; Gao, B.; Stewart, A. G.; Lee, P. V. S. On-Chip Surface Acoustic Wave and Micropipette Aspiration Techniques to Assess Cell Elastic Properties. Biomicrofluidics 2020, 14 (1), 014114. https://doi.org/10.1063/1.5138662.

(59) Rotsch, C.; Radmacher, M. Drug-Induced Changes of Cytoskeletal Structure and Mechanics in Fibroblasts: An Atomic Force Microscopy Study. Biophys. J. 2000, 78 (1), 520–535. https://doi.org/10.1016/S0006-3495(00)76614-8.

(60) Maloney, J. M.; Vliet, K. J. V. Chemoenvironmental Modulators of Fluidity in the Suspended Biological Cell. Soft Matter 2014, 10 (40), 8031–8042. https://doi.org/10.1039/C4SM00743C.

(61) Ridone, P.; Vassalli, M.; Martinac, B. Piezo1 Mechanosensitive Channels: What Are They and Why Are They Important. Biophys. Rev. 2019, 11 (5), 795–805. https://doi.org/10.1007/s12551-019-00584-5.

(62) Li, J.; Hou, B.; Tumova, S.; Muraki, K.; Bruns, A.; Ludlow, M. J.; Sedo, A.; Hyman, A. J.; McKeown, L.; Young, R. S.; Yuldasheva, N. Y.; Majeed, Y.; Wilson, L. A.; Rode, B.; Bailey, M. A.; Kim, H. R.; Fu, Z.; Carter, D. A. L.; Bilton, J.; Imrie, H.; Ajuh, P.; Dear, T. N.; Cubbon, R. M.; Kearney, M. T.; Prasad, K. R.; Evans, P. C.; Ainscough, J. F. X.; Beech, D. J. Piezo1 Integration of Vascular Architecture with Physiological Force. Nature 2014, 515 (7526), 279–282. https://doi.org/10.1038/nature13701.

(63) Zhao, Q.; Zhou, H.; Li, X.; Xiao, B. The Mechanosensitive Piezo1 Channel: A Three-Bladed Propeller-like Structure and a Lever-like Mechanogating Mechanism. FEBS J. 2019, 286 (13), 2461–2470. https://doi.org/10.1111/febs.14711.

(64) Cox, C. D.; Bavi, N.; Martinac, B. Biophysical Principles of Ion-Channel-Mediated Mechanosensory Transduction. Cell Rep. 2019, 29 (1), 1–12. https://doi.org/10.1016/j.celrep.2019.08.075.

(65) Chen, X.; Wanggou, S.; Bodalia, A.; Zhu, M.; Dong, W.; Fan, J. J.; Yin, W. C.; Min, H.-K.; Hu, M.; Draghici, D.; Dou, W.; Li, F.; Coutinho, F. J.; Whetstone, H.; Kushida, M. M.; Dirks, P. B.; Song, Y.; Hui, C.; Sun, Y.; Wang, L.-Y.; Li, X.; Huang, X. A Feedforward Mechanism Mediated by Mechanosensitive Ion Channel PIEZO1 and Tissue Mechanics Promotes Glioma Aggression. Neuron 2018, 100 (4), 799–815.e7. https://doi.org/10.1016/j.neuron.2018.09.046.

(66) Viana, N. B.; Rocha, M. S.; Mesquita, O. N.; Mazolli, A.; Maia Neto, P. A.; Nussenzveig, H. M. Towards Absolute Calibration of Optical Tweezers. Phys. Rev. E 2007, 75 (2), 021914. https://doi.org/10.1103/PhysRevE.75.021914.

(67) Schäffer, E.; Nørrelykke, S. F.; Howard, J. Surface Forces and Drag Coefficients of Microspheres near a Plane Surface Measured with Optical Tweezers. Langmuir 2007, 23 (7), 3654–3665. https://doi.org/10.1021/la0622368.

(68) Einstein, A. Investigations on the Theory of the Brownian Movement; Courier Corporation, 1956.

(69) Lenormand, G.; Millet, E.; Fabry, B.; Butler, J. P.; Fredberg, J. J. Linearity and Time-Scale Invariance of the Creep Function in Living Cells. J. R. Soc. Interface 2004, 1 (1), 91–97. https://doi.org/10.1098/rsif.2004.0010.

(70) Cox, C. D.; Bae, C.; Ziegler, L.; Hartley, S.; Nikolova-Krstevski, V.; Rohde, P. R.; Ng, C.-A.; Sachs, F.; Gottlieb, P. A.; Martinac, B. Removal of the Mechanoprotective Influence of the Cytoskeleton Reveals PIEZO1 Is Gated by Bilayer Tension. Nat. Commun. 2016, 7 (1), 10366. https://doi.org/10.1038/ncomms10366.

(71) Ho, J.; Tumkaya, T.; Aryal, S.; Choi, H.; Claridge-Chang, A. Moving beyond P Values: Data Analysis with Estimation Graphics. Nat. Methods 2019, 16 (7), 565–566. https://doi.org/10.1038/s41592-019-0470-3.

(72) Benjamin, D. J.; Berger, J. O.; Johannesson, M.; Nosek, B. A.; Wagenmakers, E.-J.; Berk, R.; Bollen, K. A.; Brembs, B.; Brown, L.; Camerer, C.; Cesarini, D.; Chambers, C. D.; Clyde, M.; Cook, T. D.; De Boeck, P.; Dienes, Z.; Dreber, A.; Easwaran, K.; Efferson, C.; Fehr, E.; Fidler, F.; Field, A. P.; Forster, M.; George, E. I.; Gonzalez, R.; Goodman, S.; Green, E.; Green, D. P.; Greenwald, A. G.; Hadfield, J. D.; Hedges, L. V.; Held, L.; Hua Ho, T.; Hoijtink, H.; Hruschka, D. J.; Imai, K.; Imbens, G.; Ioannidis, J. P. A.; Jeon, M.; Jones, J. H.; Kirchler, M.; Laibson, D.; List, J.; Little, R.; Lupia, A.; Machery, E.; Maxwell, S. E.; McCarthy, M.; Moore, D. A.; Morgan, S. L.; Munafó, M.; Nakagawa, S.; Nyhan, B.; Parker, T. H.; Pericchi, L.; Perugini, M.; Rouder, J.; Rousseau, J.; Savalei, V.; Schönbrodt, F. D.; Sellke, T.; Sinclair, B.; Tingley, D.; Van Zandt, T.; Vazire, S.; Watts, D. J.; Winship, C.; Wolpert, R. L.; Xie, Y.; Young, C.; Zinman, J.; Johnson, V. E. Redefine Statistical Significance. Nat. Hum. Behav. 2018, 2 (1), 6–10. https://doi.org/10.1038/s41562-017-0189-z.

